# Haplotype-aware inference of human chromosome abnormalities

**DOI:** 10.1101/2021.05.18.444721

**Authors:** Daniel Ariad, Stephanie M. Yan, Andrea R. Victor, Frank L. Barnes, Christo G. Zouves, Manuel Viotti, Rajiv C. McCoy

## Abstract

Extra or missing chromosomes—a phenomenon termed aneuploidy—frequently arises during human meiosis and embryonic mitosis and is the leading cause of pregnancy loss, including in the context of *in vitro* fertilization (IVF). While meiotic aneuploidies affect all cells and are deleterious, mitotic errors generate mosaicism, which may be compatible with healthy live birth. Large-scale abnormalities such as triploidy and haploidy also contribute to adverse pregnancy outcomes, but remain hidden from standard sequencing-based approaches to preimplantation genetic testing (PGT-A). The ability to reliably distinguish meiotic and mitotic aneuploidies, as well as abnormalities in genome-wide ploidy may thus prove valuable for enhancing IVF outcomes. Here, we describe a statistical method for distinguishing these forms of aneuploidy based on analysis of low-coverage whole-genome sequencing data, which is the current standard in the field. Our approach overcomes the sparse nature of the data by leveraging allele frequencies and linkage disequilibrium (LD) measured in a population reference panel. The method, which we term LD-informed PGT-A (LD-PGTA), retains high accuracy down to coverage as low as 0.05× and at higher coverage can also distinguish between meiosis I and meiosis II errors based on signatures spanning the centromeres. LD-PGTA provides fundamental insight into the origins of human chromosome abnormalities, as well as a practical tool with the potential to improve genetic testing during IVF.

**Significance Statement:** Whole chromosome gains and losses—termed aneuploidies—are the leading cause of human pregnancy loss and congenital disorders. Recent work has demonstrated that in addition to harmful meiotic aneuploidies, mitotic aneuploidies (which lead to mosaic embryos harboring cells with different numbers of chromosomes) may also be common in preimplantation embryos but potentially compatible with healthy birth. Here we developed and tested a method for distinguishing these forms of aneuploidy using genetic testing data from 8154 IVF embryos. We re-classified embryos based on signatures of meiotic and mitotic error, while also revealing lethal forms of chromosome abnormality that were hidden to existing approaches. Our method complements standard protocols for preimplantation and prenatal genetic testing, while offering insight into the biology of early development.

## Background

Whole-chromosome gains and losses (aneuploidies) are extremely common in human embryos, and are the leading causes of pregnancy loss and congenital disorders, both in the context of *in vitro* fertilization (IVF) and natural conception [1, 2]. Aneuploidy frequently arises during maternal meiosis due to mechanisms such as classical non-disjunction, premature separation of sister chromatids, and reverse segregation [3]. Such meiotic aneuploidies are strongly associated with maternal age, with risk of aneuploid conception increasing exponentially starting around age 35. Though not as well characterized, research has also demonstrated that aneuploidy of mitotic origin is prevalent during the initial post-zygotic cell divisions, potentially owing to relaxation of cell cycle checkpoints prior to embryonic genome activation [4, 5]. Such mitotic errors, which are independent of maternal or paternal age, generate mosaic embryos possessing both normal and aneuploid cells [6]. Mechanisms of mitotic aneuploidy include anaphase lag and mitotic non-disjunction [7], but also newly appreciated phenomena such as multipolar mitotic division [8, 9]. Such abnormal mitoses are surprisingly common in cleavage-stage embryos [10–12] and partially explain the high observed rates of embryonic mortality (~ 50%) during preimplantation human development.

In light of these observations, preimplantation genetic testing for aneuploidy (PGT-A) has been developed as an approach to improve IVF outcomes by prioritizing chromosomally normal (i.e., euploid) embryos for transfer, based on the inferred genetic constitution of an embryo biopsy. First introduced in the early 1990s, PGT-A has been the subject of long-standing controversy, with some meta-analyses and clinical trials drawing its benefits into question [13, 14]. Meanwhile, technical platforms underlying the test have steadily improved over time, with the current state of the art comprising low-coverage whole-genome sequencing of DNA extracted from 5-10 trophectoderm cells of blastocyst-stage embryos from days 5-7 post-fertilization [15, 16].

The improved sensitivity and resolution of sequencing-based PGT-A have placed a renewed focus on chromosomal mosaicism as a potential confounding factor for diagnosis and interpretation [17]. Mosaicism within a PGT-A biopsy may generate a copy number profile that is intermediate between euploid and aneuploid expectations, though it is important to note that technical artifacts can mimic such signatures [18]. At the whole-embryo scale, mosaic aneuploidies may be prevalent but systematically underestimated due to the reliance of PGT-A on biopsies of one or few cells, which by chance may fail to sample low-level aneuploid lineages [19]. While uniform aneuploidies arising from meiotic errors are unambiguously harmful, mosaic aneuploidies are potentially compatible with healthy live birth [20–22]. Research using mouse models and human gastruloids has recently offered initial insights into the selective elimination of aneuploid cells from mosaic embryo via lineagespecific mechanisms of apoptosis [23–25].

Another underappreciated challenge in the analysis of sequencing-based PGT-A data is the detection of complex abnormalities including errors in genome-wide ploidy (e.g., triploidy and haploidy). Because existing algorithms detect chromosome abnormalities by comparing the normalized counts of aligned sequencing reads (i.e., depth of coverage) across chromosomes within a sample, inferences are compromised when many or all chromosomes are affected. Extreme cases such as haploidy and triploidy may evade detection entirely and be falsely interpreted as normal euploid samples. Such classification errors are particularly concerning given the association of ploidy abnormalities with molar pregnancy and miscarriage [26, 27]. Triploidy in particular comprises more than 10% of cytogenetically abnormal miscarriages [28].

The ability to reliably distinguish meiotic- and mitotic-origin aneuploidies, as well as complex and genomewide errors of ploidy, may thus prove valuable for enhancing IVF outcomes. Notably, trisomy (and triploidy) of meiotic origin is expected to exhibit a unique genetic signature, characterized by the presence of three distinct parental haplotypes (i.e., “both parental homologs” [BPH] from a single parent) in contrast with the mitotic trisomy signature of only two distinct haplotypes chromosome-wide (i.e., “single parental homolog” [SPH] from each parent; Fig. 1) [29]. Conversely, monosomy and haploidy will exhibit genetic signatures of only a single haplotype chromosome- or genome-wide, respectively. To date, few methods have explicitly attempted to use these signatures to distinguish these forms of aneuploidy. Exceptions include approaches that require genetic material from both parents and embryo biopsies [30–33] as well as targeted sequencing approaches that require alternative methods of library preparation to sequence short amplicons to higher coverage [34].

**Figure 1.**
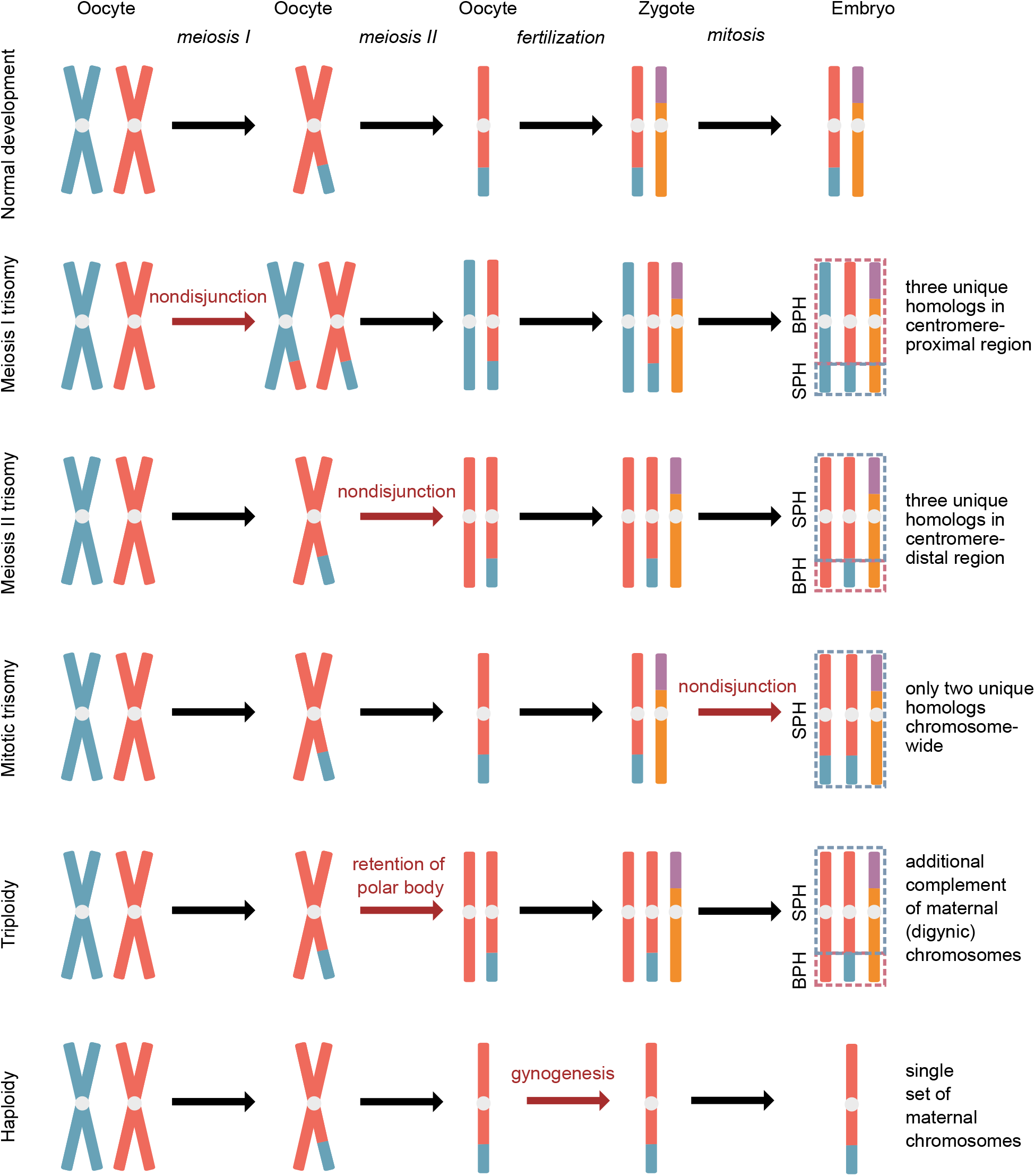
Signatures of various forms of chromosome abnormality with respect to their composition of identical and distinct parental homologs. Normal gametogenesis produces two genetically distinct copies of each chromosome—one copy from each parent—that comprise mosaics of two homologs possessed by each parent. Meiotic-origin trisomies may be diagnosed by the presence of one or more tracts with three distinct parental homologs (i.e., transmission of both parental homologs [BPH] from a given parent). In contrast, mitotic-origin trisomies are expected to exhibit only two genetically distinct parental homologs chromosome-wide (i.e., duplication of a single parental homolog [SPH] from a given parent). Triploidy and haploidy will mirror patterns observed for individual meiotic trisomies and monosomies, respectively, but across all 23 chromosome pairs—a pattern that confounds standard coverage-based analysis of PGT-A data.

Here we describe a statistical approach to classify aneuploidies using genotype information encoded in low-coverage whole-genome sequencing data from PGT-A. Inspired by the related challenge of imputation [35–40], our method overcomes the sparse nature of the data by leveraging information from a population reference panel. We test our method on simulated and empirical data of varying sequencing depths, meiotic recombination patterns, and patient ancestries, evaluating its strengths and limitations under realistic scenarios. At higher coverage, we further demonstrate that our method permits the mapping of meiotic crossovers on trisomic chromosomes, as well as the distinction of trisomies originating in meiosis I and meiosis II. Our method reveals new biological insight into the fidelity of meiosis and mitosis, while also holding promise for improving preimplantation genetic testing.

## Results

### Genotypic signatures of meiotic and mitotic aneuploidy

Most contemporary implementations of PGT-A are based on low-coverage (<0.05x) whole-genome sequencing of 5-10 trophectoderm cells biopsied from blastocyst-stage embryos at day 5, 6, or 7 postfertilization [15, 16]. Given the level of coverage and the paucity of heterozygous sites within human genomes, standard approaches for analyzing such data typically ignore genotype information and instead infer aneuploidies based on deviations in relative coverage across chromosomes within a sample. Inspired by genotype imputation [35–40] and related forensic methods [41], we hypothesized that even low-coverage data could provide orthogonal evidence of chromosome abnormalities, complementing and refining the inferences obtained from coverage-based methods. As in imputation, the information content of the genotype data is greater than first appears by virtue of linkage disequilibrium (LD)— the population genetic correlation of alleles at two or more loci in genomic proximity, which together comprise a haplotype [42]. We developed an approach to quantify the probability that two sequencing reads originated from the same chromosome, based on known patterns of LD among alleles observed on those reads. Hereafter, we refer to our method as LD-informed PGT-A (LD-PGTA).

Specifically, in the case of trisomy, we sought to identify a signature of meiotic error wherein portions of the trisomic chromosome are composed of two distinct homologs from one parent and a third distinct homolog from the other parent (BPH; Fig. 1). In contrast, trisomies arising via post-zygotic mitotic errors are composed of two identical copies of one parental homolog and a second distinct homolog from the other parent (SPH). Although this chromosome-wide SPH signature may also capture rare meiotic errors with no recombination (Fig. 1), BPH serves as an unambiguous signature of deleterious meiotic aneuploidy. In the presence of recombination, meiotic trisomies will comprise a mixture of BPH and SPH tracts. BPH tracts that span the centromere are consistent with mis-segregation of homologous chromosomes during meiosis I, while BPH tracts that lie distal to the centromere are consistent with missegregation of sister chromatids during meiosis II [43].

### LD-PGTA: a statistical model to classify aneuploidies

Distinguishing these scenarios in low-coverage data (< 1×) is challenging due to the fact that reads rarely overlap one another at sites that would distinguish the different homologs. Our classifier therefore uses patterns of allele frequencies and LD from large population reference panels to account for potential relationships among the sparse reads. Specifically, the length of the trisomic chromosome is divided into genomic windows, on a scale consistent with the length of typical human haplotypes (10^4^ - 10^5^ bp). For each genomic window, several reads are sampled, and the probabilities of the observed alleles under the BPH trisomy and SPH trisomy hypotheses (i.e., likelihoods) are computed.

Our likelihood functions are based on the premise that the probability of drawing reads from the same haplotype differs under different ploidy hypotheses, which are defined by various configurations of genetically distinct or identical homologs (Fig. 1). If a pair of reads originates from two identical homologs, the probability of observing the alleles on these reads is given by the joint frequency of the linked alleles. On the other hand, if a pair of reads originates from two different homologs, the probability of observing their associated alleles is simply equal to the product of the allele frequencies, since the alleles are unlinked. Because the BPH and SPH hypotheses are defined by different ratios of identical and distinct homologs, the probability of sampling reads from the same homolog also differs under each hypothesis (1/3 and 5/9, respectively; Fig. 1).

Allele frequencies and joint allele frequencies (i.e., haplotype frequencies) are in turn estimated through examination of a phased population genetic reference panel, ideally matched to the ancestry of the target sample. Such estimates have the virtue that they do not depend on theoretical assumptions, but simply on the sample having been randomly drawn from the genetically similar populations. A drawback, which we take into account, is that reliable estimates of the frequencies of rare alleles or haplotypes require large reference panels. In practice, it is sufficient to construct the reference panels using statistically phased genotypes from large surveys of human genetic diversity, such as the 1000 Genomes Project *(n* = 2504 individuals).

Given the importance of admixture in contemporary populations, we also generalized our models to allow for scenarios involving admixture in the parental generation, as well as more distant and complex admixture scenarios (see Methods). These admixture-aware models only require knowledge of the ancestry of the target sample (i.e., embryo), which has a practical advantage for PGT-A where parental genotype data is typically unavailable.

Within each genomic window, we compare the likelihoods under the BPH and SPH hypotheses by computing a log-likelihood ratio (LLR) and use the *m* out of *n* bootstrap procedure [44] to assess uncertainty (see Methods; Fig. 2). LLRs are then aggregated across consecutive genomic windows comprising larger intervals (i.e., “bins”). The mean and variance of the combined LLR is used to compute a confidence interval for each bin, which can then be classified as supporting the BPH or SPH hypothesis, depending on whether the bounds of the confidence interval are positive or negative, respectively. Confidence intervals that span zero remain inconclusive, and are classified as “ambiguous”.

**Figure 2.**
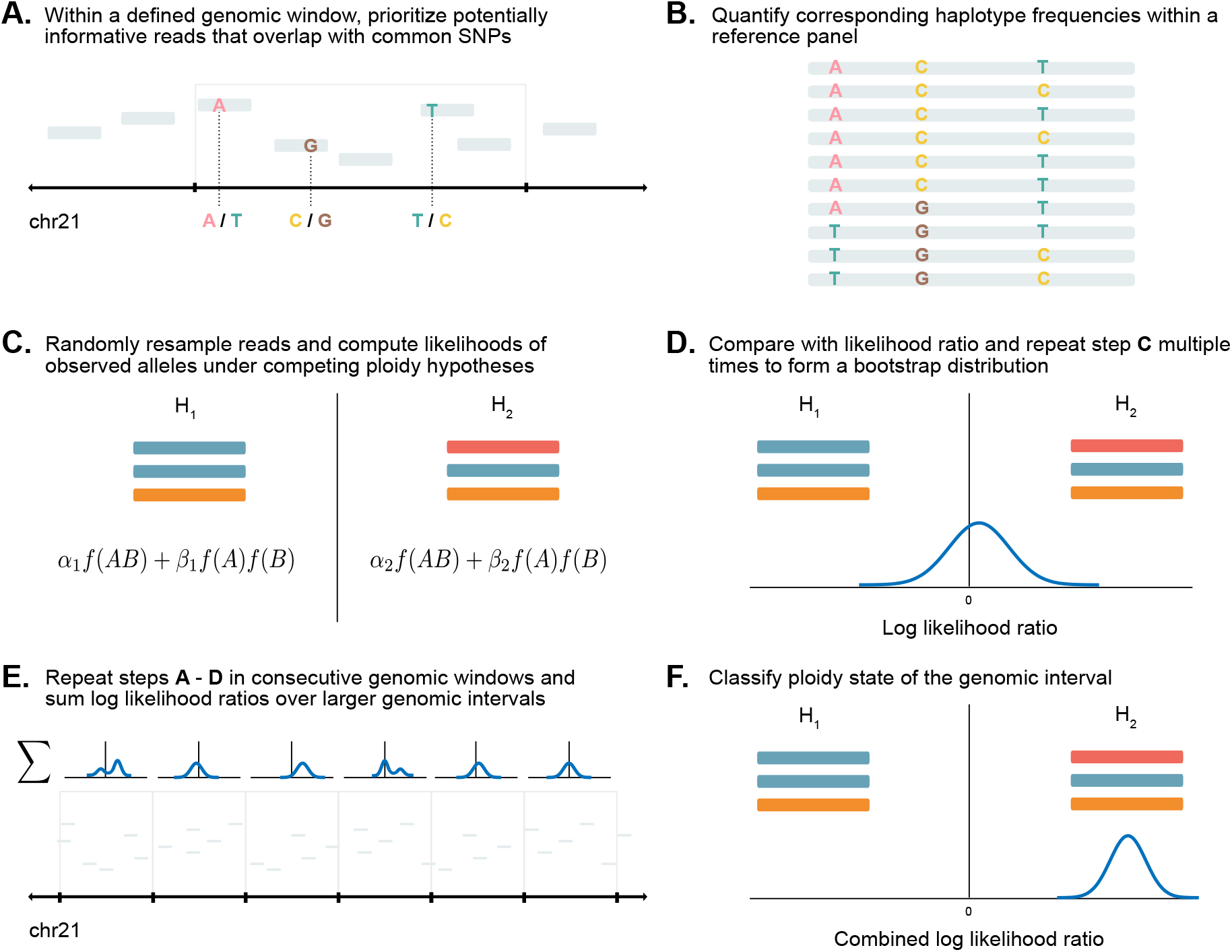
A. Within defined genomic windows, select reads overlapping informative SNPs that tag common haplotype variation. B. Obtain joint frequencies of corresponding SNPs from a phased panel of ancestry-matched reference haplotypes. C. Randomly resample 2-18 reads and compute probabilities of observed alleles under two competing ploidy hypotheses. D. Compare the hypotheses by computing a likelihood ratio and estimate the mean and variance by re-sampling random sets of reads using a bootstrapping approach. E. Repeat steps A - D for consecutive non-overlapping genomic windows and aggregate the log likelihood ratios over larger genomic intervals. F. Use the mean and variance of the combined log likelihood ratio to estimate a confidence interval and classify the ploidy state of the genomic interval.

### Benchmarking LD-PGTA with simulated sequences

To evaluate our method, we simulated sequencing reads from BPH and SPH trisomies according to their defining haplotype configurations (Fig. 1). Our simulations assumed uniform depth of coverage, random mating (by randomly drawing haplotypes from the 1000 Genomes Project), and equal probability of drawing a read from any of the homologs. We varied the mean depths of coverage (0.01x, 0.05x, and 0.1x) and read lengths (36-250 bp), testing model assumptions and performance over a range of plausible scenarios.

In order to benchmark and optimize LD-PGTA, we developed a generalization of the ROC curve [45] for scenarios that include an ambiguous class (i.e., bins with a confidence interval spanning zero), which we hereafter denote as the “balanced” ROC curve. For a given discrimination threshold, a balanced true (false) positive rate, denoted as BTPR (BFPR), is defined as the average of the true (false) positive rate of predicting BPH and SPH trisomy (Figs. 3a and 3b). The balanced ROC curve thus depicts the relationship between the BTPR and BFPR at various confidence thresholds.

**Figure 3.**
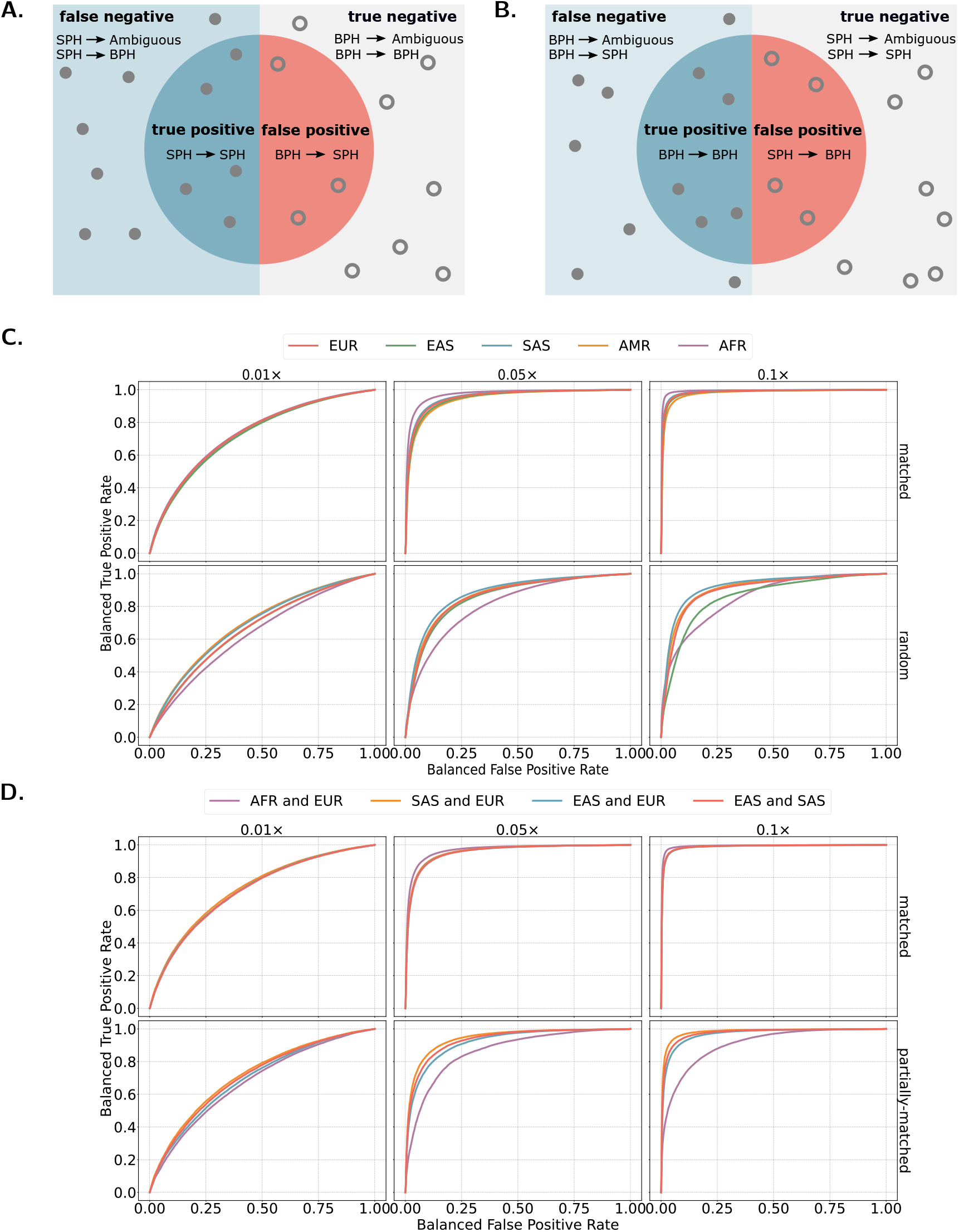
The upper left (right) diagram depicts relative trade-offs between instances classified correctly as SPH (BPH) and instances misclassified as SPH (BPH). For example, all instances of actual class SPH that were classified as instances of predicated class BPH are denoted by SPH ⊒ BPH. The balanced true (false) positive rate is defined as an average of the true (false) positives rates from both diagrams (Eqs. (13) and (14)). C. Balanced ROC curves for BPH vs. SPH with matched and random reference panels for non-admixed embryos, varying depths of coverage. D. Balanced ROC curves for BPH vs. SPH with matched and partially-matched reference panels for recent admixed embryos, varying depths of coverage. Here the randomly mismatched reference panel corresponds to either the paternal or maternal ancestry. Each balanced ROC curve reflects an average over bins across the genome. We averaged both the BTPR and BFPR for common *z*-scores across bins.

We generated balanced ROC curves for bins within simulated trisomic chromosomes (BPH and SPH) for samples of non-admixed ancestry and a correctly specified reference panel (Fig. 3c). At depths of 0.01x, 0.05x and 0.1x, LD-PGTA achieved a mean BTPR across ancestry groups of 33.4%, 88.6% and 97.5%, respectively, with a BFPR of 10%. Expanding our view across all classification thresholds, we found that the area under the balanced ROC curves was 0.72, 0.95 and 0.99 at depths of 0.01x, 0.05x and 0.1x, respectively. Our results thus demonstrate that as the depth of coverage increases from 0.01x to 0.1x, the balanced ROC curve approaches a step function, nearing ideal classification performance.

We also observed that short read lengths performed better than longer reads at low coverage (Fig. S1). While potentially counter-intuitive, this relationship arises due to the fact that coverage is a linear function of the read length as well as the number of reads. Thus, holding coverage constant, a large number of shorter reads will achieve more uniform coverage than a small number of longer reads.

For meiotic-origin trisomies, meiotic crossovers should manifest as switches between tracts of BPH and SPH trisomy. To test the utility of our method for revealing such recombination events, we simulated trisomic chromosomes with random mixtures of BPH and SPH tracts, while varying the sequencing depth, as well as the chromosome length and size of the corresponding genomic windows (Fig. 4). Applying LD-PGTA to the simulated data, we examined the resulting changes in the LLRs of consecutive bins and their relation to simulated meiotic crossovers. While muted signatures could be discerned at the lowest coverage of 0.01 ×, the signatures were difficult to distinguish from background noise and spatial resolution was poor. In contrast, crossovers were more pronounced at higher sequencing depths of 0.05× and 0.1 × and closely corresponded to simulated crossover breakpoints. For certain chromosomes (e.g., Chr11, Chr13, and Chr18) and at the higher sequencing depths, the LLRs observed in centromere-flanking bins provided an indication of the originating stage of the trisomy (here simulated as meiosis II errors). We note, however, that the bin encompassing the centromere itself was rarely informative, as such genomic regions are typically composed of repetitive heterochromatin that is inaccessible to standard short-read mapping and genotyping methods.

**Figure 4.**
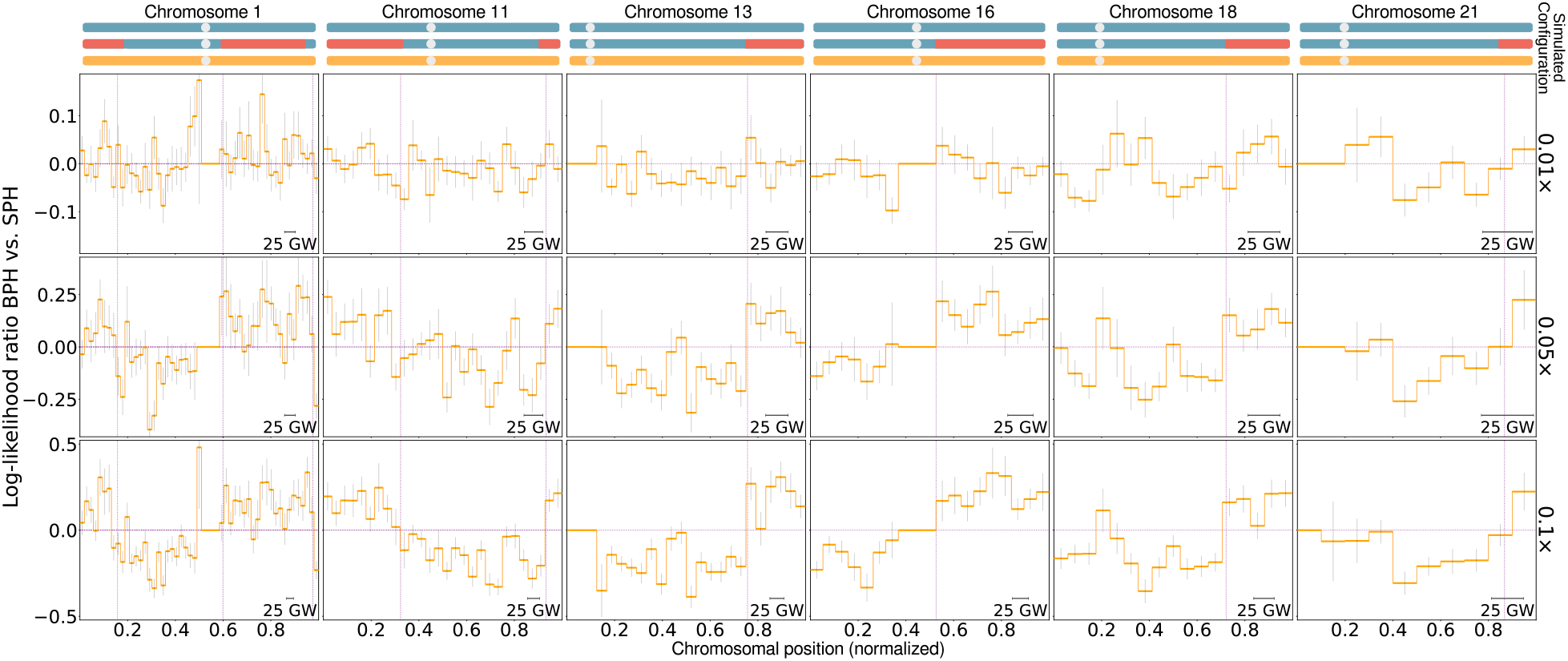
Demonstration of the detection of meiotic crossovers from low-coverage PGT-A data. Trisomies were simulated with varying locations of meiotic crossovers, as depicted in the upper diagrams and varying depths of coverage (0.01x, 0.05×, and 0.1x). Confidence intervals correspond to a *z*-score of one (confidence level of 68.3%). The size of the genomic windows varies with the coverage, while the size of the bins is kept constant.

### Evaluating sensitivity to ancestries of the target sample and reference panel

We next extended our simulations to test the sensitivity of our classifier to specification of a reference panel that is appropriately matched to the ancestry of the target sample. Specifically, we simulated samples by drawing haplotypes from a reference panel composed of one of the five super-populations of the 1000 Genomes Project, then applied LD-PGTA while either specifying a reference panel composed of haplotypes from the same super-population (i.e., “matched”) or from a mixture of all super-populations (i.e., “mismatched”). Notably, any phased genomic dataset of matched ancestry could be used in place of the 1000 Genomes Project dataset, with performance maximized in large datasets with closely matched ancestries.

While classification performance was high across all continental ancestry groups under the matched-ancestry scenario, performance moderately declined in all scenarios where the reference panel was misspecified (Fig. 3c; Figs. S3a, S4a and S5a). Due to such misspecification, the mean BTPR across ancestry groups declined by 8.5%, 28.3% and 26.4%, at a BFPR of 10% and depths of 0.01 ×, 0.05× and 0.1 ×, respectively. Moreover, the area under the balanced ROC curves decreased by 0.06, 0.10 and 0.10 at depths of 0.01×, 0.05× and 0.1×, respectively. This performance decline mimics that of related methods such as polygenic scores and arises by consequence of systematic differences in allele frequencies and LD structure between the respective populations [46–48]. As an example of such effects, we observed that using an African reference panel for a European target sample produced a bias in favor of SPH trisomy (Fig. S5). Conversely, using a European reference panel for an African target sample produced a bias in favor of BPH trisomy.

Seeking to test classification performance on individuals of admixed ancestries, we extended our simulation procedure to generate test samples composed of chromosomes drawn from distinct super-populations. In following with our previous analysis, we tested our method with and without correct specification of the component ancestries contributing to the admixture. Specifically, we implemented the latter scenario by specifying a single reference panel that matched the ancestry of eitherthe maternal or paternal haplotypes (i.e., “partially-matched”; Fig. 3d). Our results demonstrated that the performance of LD-PGTA for correctly specified admixture scenarios was comparable to that observed for non-admixed scenarios. The ratio of mean AUC across ancestries and coverages for non-admixed samples versus admixed samples was 1.0. The impacts of reference panel misspecification were again greatest for admixed individuals with African ancestry, likely reflecting differences in structure of LD (shorter haplotypes) among African populations. For the African-European admixture scenario the BTPR decreased by 5.1%, 30.9% and 28.7% at a BFPR of 10% and depths of 0.01×, 0.05× and 0.1× when admixture was misspecified.

### Application to PGT-A data

We next proceeded to apply our method to existing data from PGT-A, thereby further evaluating its performance and potential utility under realistic clinical scenarios. Data were obtained from IVF cases occurring between 2015 and 2020 at the Zouves Fertility Center (Foster City, USA). Embryos underwent trophectoderm biopsy at day 5, 6, or 7 post-fertilization, followed by PGT-A using the Illumina VeriSeq PGS kit and protocol, which entails sequencing on the Illumina MiSeq platform (36 bp single-end reads). The dataset comprised a total of 8154 embryo biopsies from 1640 IVF cases, with maternal age ranging from 22 to 56 years (median = 38). Embryos were sequenced to a median coverage 0.0083x per sample, which we note lies near the lower limit of LD-PGTA and corresponds to an expected false positive rate of ~ 0.02 at a 50% confidence threshold for classification of chromosome-scale patterns of SPH and BPH trisomy Fig. S7a.

### Ancestry inference for reference panel selection

Given the aforementioned importance of the ancestry of the reference panel, we used LASER [49, 50] to perform automated ancestry inference of each embryo sample from the low-coverage sequencing data. LASER applies principal components analysis (PCA) to genotypes of reference individuals with known ancestry. It then projects target samples onto the reference PCA space, using a Procrustes analysis to overcome the sparse nature of the data. Ancestry of each target sample is then deduced using a k-nearest neighbors approach.

Our analysis revealed a diverse patient cohort, consistent with the demographic composition of the local population (Fig. 5). Specifically, we inferred a total of 2037 (25.0%) embryos of predominantly East Asian ancestries, 900 (11.0%) of South Asian ancestries, 2554 (31.3%) of European ancestries, 669 (8.2%) of admixed American ancestries, and 44 (0.5%) of African ancestries, according to the reference panels defined by the 1000 Genomes Project. Interestingly, we also observed 1936 (23.7%) embryos with principal component scores indicating admixture between parents of ancestries from one of the aforementioned super-populations, which we thus evaluated using the generalization of PGT-A for admixed samples. More specifically, we inferred a total of 1264 (15.5%) embryos to possess admixture of East Asian and European ancestries, 447 (5.5%) of South Asian and European ancestries, 129 (1.6%) of East and South Asian ancestries and 96 (1.2%) of African and European ancestries. The ancestry of 14 (0.2%) embryos remained undetermined. To further test the robustness of these ancestry inferences with low amounts of input data, we separately analyzed Chromosome 1 and compared to inferences for the rest of the genome. Even when restricting to this small proportion of the genome (< 10%), we observed concordances of 87.1% and 81.9% for putative non-admixed and admixed individuals, respectively.

**Figure 5.**
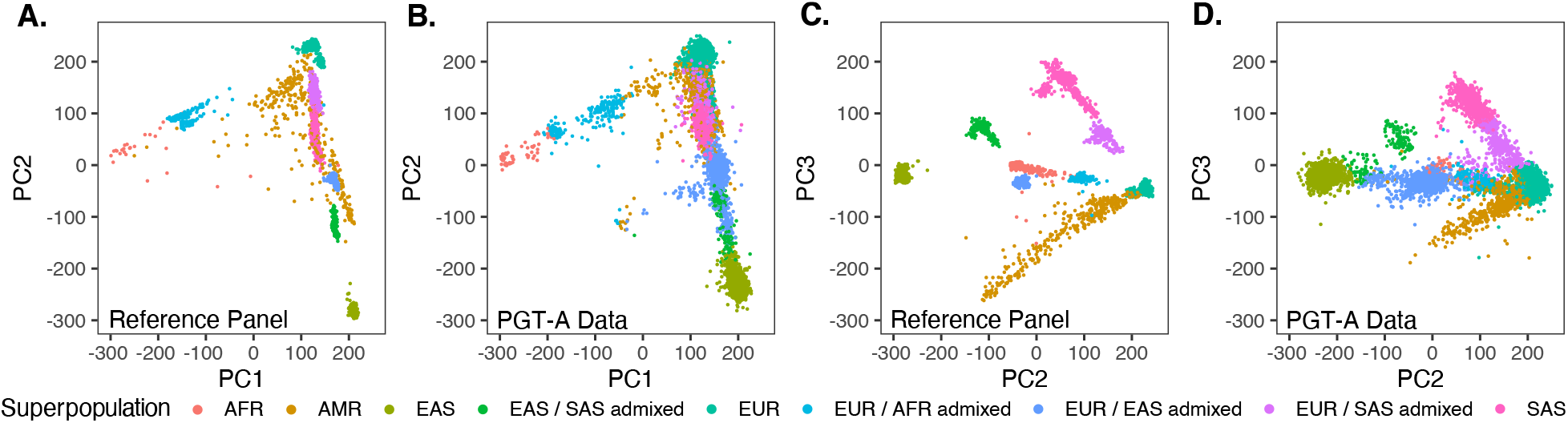
Ancestry inference from low-coverage PGT-A data informs the selection of matched reference panels. Principal component axes were defined based on analysis of 1000 Genomes reference samples (panels A and C), and colored according to superpopulation annotations. Low-coverage embryo samples (genome-wide excluding chromosome 1) were then projected onto these axes using a Procrustes approach implemented with LASER (v2.0) [50] (panels B and D) and their ancestries were classified using k-nearest neighbors of the first four principal component axes (*k* = 10). Projections of first-generation admixed reference samples were approximated as the mean of random samples from each of the component superpopulations. Panels A and B depict principal components 1 and 2, while panels C and D depict principal components 2 and 3, together capturing continental-scale patterns of global ancestry.

### Preliminary analysis with a coverage-based classifier

Ploidy status of each chromosome from each embryo biopsy was first inferred using the Illumina BlueFuse Multi Software suite in accordance with the VeriSeq protocol. Similar in principle to several open source tools [51], BlueFuse Multi detects aneuploidies based on the coverage of reads aligned within 2500 bins that are distributed along the genome, normalizing and adjusting for local variability in GC content and other potential biases. It then reports the copy number estimates for each bin (and interpolates between bins), as well as the copy number classification of each full chromosome (as a “gain”, “loss”, or neither) and numerous auxiliary metrics to assist with quality control.

Of the original 8154 embryos, we excluded 138 embryos with low signal-to-noise ratios (Derivative Log Ratio [DLR] ≥ 0.4), as well as an additional 970 embryos with low coverages that were unsuitable for analysis with LD-PGTA (mean coverage < 0.005×). This resulted in a total of 7046 embryos used for all downstream analyses. We focused our analysis on aneuploidies affecting entire chromosomes (i.e., non-segmental aneuploidies; see Methods), as these are the hypotheses that LD-PGTA is designed to test at an extremely low coverage. (Fig. 1). Aggregating the BlueFuse Multi results from these embryos, we identified 2006 embryos that possessed one or more non-mosaic whole-chromosome aneuploidy (28.5%), including 869 (12.3%) with one or more chromosome gain and 1186 (16.8%) with one or more chromosome loss. Of these, a total of 494 embryos (7.0%) possessed aneuploidies involving two or more chromosomes. A total of 307 (4.4%) embryos possessed one or more chromosomes in the mosaic range, including 94 (1.3%) embryos that also harbored non-mosaic aneuploidies. The rate of non-mosaic aneuploidy was significantly associated with maternal age (Quasibinomial GLM: 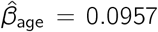, *SE* = 0.0080, *P* < 1 × 10^−10^), while the rate of mosaic aneuploidy was not (Quasibinomial GLM: 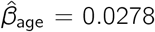, *SE* = 0.0143, *P* = 0.0517) replicating well described patterns from the literature [1]. On an individual chromosome basis, the above results correspond to a total of 1657 whole-chromosome losses (1456 complete and 201 in the mosaic range) and 1331 whole-chromosome gains (1063 complete and 268 in the mosaic range) (Fig. 6a). These conventional coverage-based results served as the starting point for refinement and sub-classification with LD-PGTA.

**Figure 6.**
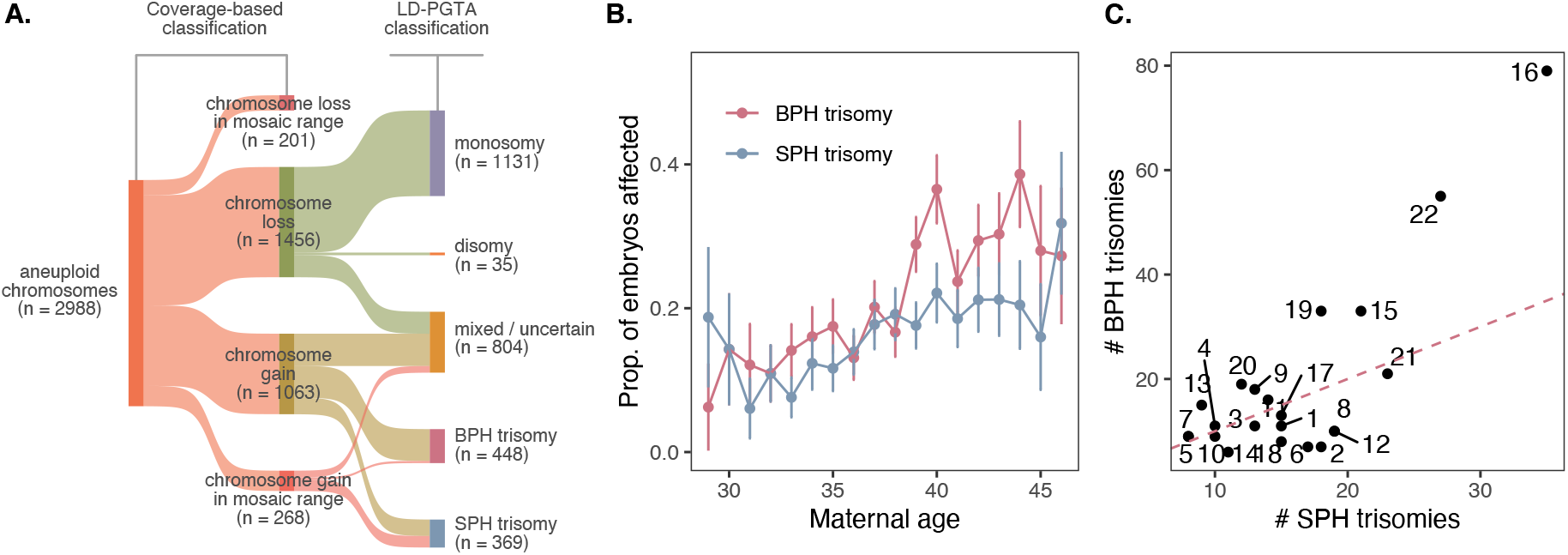
Application of LD-PGTA to low-coverage sequencing-based data. A. LD-PGTA was used to refine classification results from 2988 chromosomes originally diagnosed as whole-chromosome (i.e., non-segmental) aneuploid based on standard coverage-based analysis with Bluefuse Multi, including strong validation of putative monosomies, as well as subclassification of BPH and SPH trisomies. B. Association of BPH and SPH trisomies with maternal age. C. LD-PGTA classifications of BPH versus SPH trisomy, stratifying by individual chromosome. BPH trisomy is strongly enriched on chromosomes 16 and 22, while signatures of SPH trisomy are more evenly distributed among the various autosomes. Chromosomes were classified using a 50% confidence interval.

### Sub-classification of meiotic and mitotic trisomies

LD-PGTA is intended to complement coverage-based methods, using orthogonal evidence from genotype data to support, refute, or refine initial ploidy classifications. The relevant hypotheses for LD-PGTA to test are therefore based on the initial coverage-based classification. Applying LD-PGTA to the 1456 putative non-mosaic whole-chromosome losses, we confirmed the monosomy diagnosis for 1131 (77.7%) chromosomes, designated 290 as ambiguous (i.e., 50% confidence interval spanning 0), and obtained conflicting evidence favoring disomy for only 35 chromosomes (2.4%). We note that the latter group is within the expected margin of error (FPR = 5.1%) if all 1456 chromosomes were in fact monosomic, given the selected confidence threshold of 50%.

Seeking to better understand the molecular origins of trisomy, we next applied LD-PGTA to the 1331 originally diagnosed whole-chromosome gains (1063 complete and 268 in the mosaic range). Of the non-mosaic gains, a total of 420 (39.5%) chromosomes exhibiting evidence of BPH trisomy (i.e., meiotic origin), 217 (20.4%) chromosomes exhibiting evidence of full SPH trisomy (i.e., mitotic origin or meiotic origin lacking recombination), and 426 (40.1%) chromosomes exhibiting evidence of a mixture of BPH and SPH tracts (i.e., meiotic origin with recombination) or uncertain results. In contrast, the vast majority of the 268 putative mosaic chromosome gains exhibited evidence of SPH trisomy (152 chromosomes [56.7%]), while only 28 (10.4%) exhibited evidence of BPH trisomy and 88 (32.8%) remained ambiguous. We observed that BPH trisomies exhibited a nominally stronger association with maternal age than SPH trisomies, consistent with previous literature demonstrating a maternal age effect on meiotic but not mitotic aneuploidy, though the difference was not statistically significant (Quasibinomial GLM: 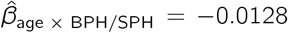, *SE* = 0.0184, *P* = 0.486; Fig. 6b) [29]. Intriguingly, Chromosomes 16 (binomial test: *P* = 5.3 x 10^−6^) and 22 (binomial test: *P* = 2.7 x 10^−7^) were strongly enriched for BPH versus SPH trisomies, again consistent with a known predisposition of these two chromosomes to missegregation during maternal meiosis, including based on their strong maternal age effect (Fig. 6c) [29, 52].

Scanning along consecutive bins across a trisomic chromosome, switches between tracts of BPH and SPH trisomy indicate the occurrence of recombination, offering a method for mapping meiotic crossovers and studying their role in the genesis of aneuploidy. To detect these crossovers in practice, we omitted all ambiguous bins and then approximated each crossover breakpoint as the mid-point between the centers of nearest-neighbor bins, where one was unambiguously classified as BPH and the other as SPH. Although the low coverage nature of this particular dataset offered very coarse resolution (~5 Mbp; see Discussion), we nevertheless identified 1614 putative meiotic crossovers and mapped their heterogeneous distribution across 2149 trisomic chromosomes (conditioning on coverage >0.01 x and based on a ±1 SD confidence threshold; Fig. 7).

**Figure 7.**
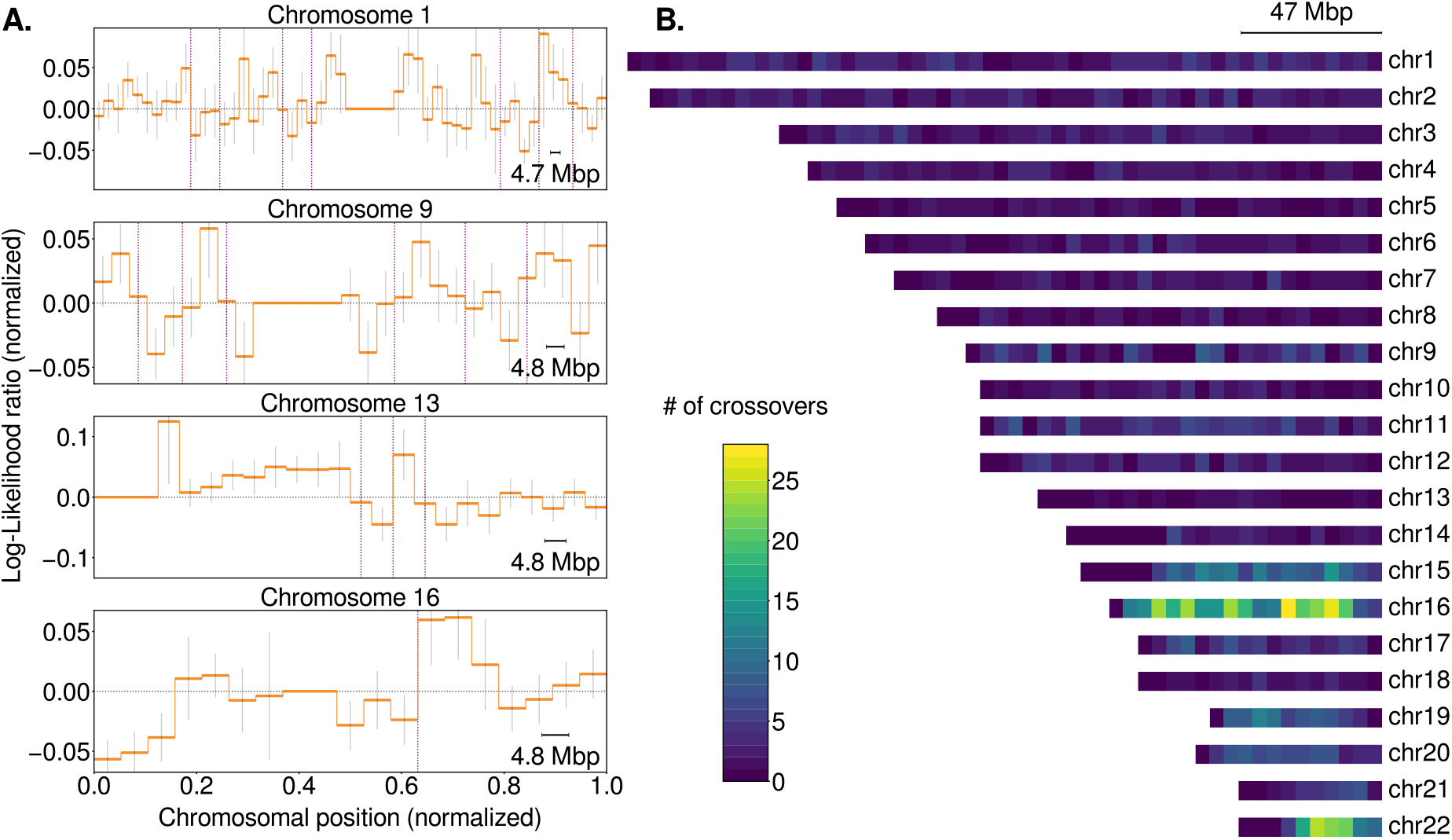
Mapping of meiotic crossovers on putative trisomic chromosomes based on inferred switches between tracts of BPH and SPH trisomy. Samples with less than 0.01x coverage were excluded from the analysis. A. Putative crossovers (vertical lines) observed for trisomic chromosomes of four representative samples. Confidence intervals correspond to a *z*-score of one. B. Annotated ideogram of meiotic crossovers detected in each chromosomal bin of all autosomes across the entire sample.

### Hidden abnormalities in genome-wide ploidy

Because of their reliance on differences in normalized coverage of reads aligned to each chromosome, sequencing-based implementations of PGT-A are typically blind to haploidy, triploidy, and other potential genome-wide aberrations. While differences in coverage between the X and Y chromosome may offer a clue about certain forms of triploidy [53], this scenario does not apply to haploidy or to forms of triploidy where the Y chromosome is absent. Moreover, even when present, the short length of the Y chromosome diminishes the ratio of signal to noise and limits its diagnostic utility. As such, haploid and triploid embryos (especially those lacking Y chromosomes) are routinely mis-classified as diploid by coverage-based methods for PGT-A analysis [34].

In order to overcome these challenges, we extended LD-PGTA to detect abnormalities in genome-wide ploidy, effectively combining evidence for or against specific chromosome abnormalities across the entire genome. In the case of haploidy, this entailed aggregating the LLRs of the comparison between disomy and monosomy across all genomic bins, while in the case of triploidy, we aggregated the LLRs of the comparison between disomy and BPH trisomy across all genomic bins. Because the chromosomes of triploid samples will possess tracts of both BPH and SPH trisomy, our power for detection is lower than for haploidy, and it thus requires a less stringent threshold (see Methods; Fig. S8a).

We started our analysis with the the 4495 embryos that were initially classified as euploid based on conventional coverage-based tests (i.e., Bluefuse Multi). To take a more conservative approach, we applied additional filtering to consider only embryos with >0.06x and >0.03x with at least 1000 and 250 informative genomic windows for the classification of triploidy and haploidy, respectively. In the case of triploidy, this confined the initial pool of embryos to 3821, among which we identified 12 (0.31%) as likely triploid (see Methods; Eq. (25); Fig. 8). In the case of haploidy, our stringent filtering criteria restricted our analysis to an initial pool of 1320 embryos, among which we identified 11 (0.83%) as likely haploid (see Methods; Eq. (11); Fig. 8). Of the 12 putative triploid embryos, 5 (42%) exhibited an XXX composition of sex chromosomes. In contrast, all of the haploid embryos possessed an X chromosome and no Y chromosome, consistent with previous studies of IVF embryos that suggested predominantly gynogenetic origins of haploidy, both in the context of intracytoplasmic sperm injection (ICSI) and conventional IVF [54, 55]. Among the 12 putative triploid embryos, we observed 5 embryos with coverage-based signatures of XXY sex chromosome complements. Moreover, 6 of the 12 embryos which we classified as haploid were independently noted as possessing only a single pronucleus (i.e., monopronuclear or 1PN) at the zygote stage upon retrospective review of notes from the embryology lab. Importantly, however, microscopic assessment of pronuclei is known to be prone to error and is imperfectly associated with ploidy status and developmental potential [56]. Nevertheless, these results offer independent validation of our statistical approach, which may be used to flag diploid-testing embryos as harboring potential abnormalities in genome-wide ploidy.

**Figure 8.**
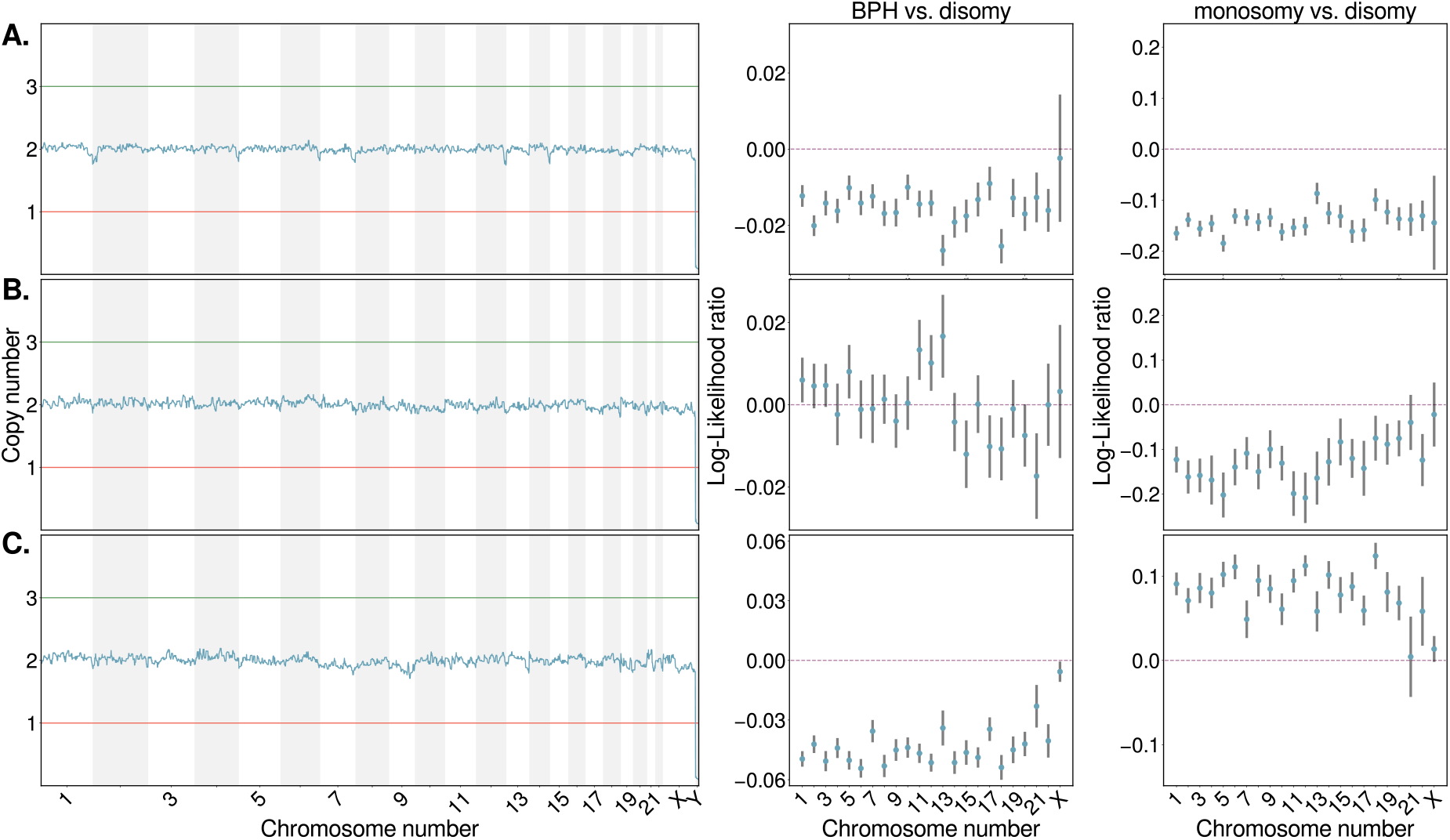
Visualization of results from representative putative diploid (A.), triploid (B.), and haploid (C.) samples. Copy number estimates obtained using a standard coverage-based approach (BlueFuse Multi) are depicted in the left column and are indicative of diploidy. LD-PGTA results are depicted in the right two columns and suggest genome-wide abnormalities in ploidy for the latter two samples.

## Discussion

Aneuploidy affects more than half of human embryos and is the leading cause of pregnancy loss and IVF failure. PGT-A seeks to improve IVF outcomes by prioritizing euploid embryos for transfer. Current low-coverage sequencing-based implementations of PGT-A rely entirely on comparisons of the depth of coverage of reads aligned to each chromosome. In such low coverage settings, genotype observations are so sparse that they are typically regarded as uninformative.

Here we showed that by leveraging patterns of LD, even sparse genotype data are sufficient to capture signatures of aneuploidy, especially when aggregating over large genomic intervals such as entire chromosomes. In doing so, our method, termed LD-PGTA, reveals additional information that is hidden to standard coverage-based analyses of PGT-A data, including the designation of meiotic and mitotic trisomies, as well as the discovery of errors in genome-wide ploidy (i.e., haploidy and triploidy). Using simulations, we demonstrated the high accuracy of LD-PGTA even at the extremely low depths of coverage (0.05x). We then showcased the utility of our method through application to an existing PGT-A dataset composed of 7046 IVF embryos, stratifying trisomies of putative meiotic and mitotic origin, and reclassifying 11 presumed diploid embryos as haploid and an additional 12 as triploid.

The key innovation of LD-PGTA is the use of allele frequencies and patterns of LD as measured in an external reference panel such as the 1000 Genomes Project [57]. Because these parameters vary across human populations, the accuracy of our method depends on the correct specification of the reference panel with respect to the ancestry of the target sample. This challenge is analogous to that described in several recent studies investigating the portability of polygenic scores between populations, which are similarly biased by population differences in allele frequencies and patterns of LD [46–48]. In contrast to polygenic scores, however, our simulations suggest that the practical effects of reference panel misspecification on aneuploidy classification are typically modest. Moreover, our analysis of PGT-A data demonstrates that even at very low depths of coverage, existing extensions of principal components analysis [50] are sufficient to infer sample ancestries and guide the selection of an appropriate matched reference panel. However, this still depends on the availability of those genetic reference panels. While studies such as the 1000 Genomes Project [57] offer a broad sampling of global populations, certain populations remain underrepresented (including many African populations, as well as indigenous populations around the world). We anticipate that the performance of our method will improve as genomic databases grow larger and more diverse.

While LD-PGTA is conceptually related to genotype imputation in the sense that both exploit patterns of LD from a population reference panel, the goals of these two tasks are distinct. Whereas genotype imputation assumes some ploidy state and seeks to infer missing genotypes (e.g., for SNPs that are not assayed by a genotyping microarray) [36, 38–40], LD-PGTA uses the haplotype patterns to quantify evidence of different ploidy configurations. By combining evidence over large genomic intervals, LD-PGTA achieves classification power at coverages so low (<0.05x) that accurate genotype imputation is not possible [39]. One intriguing area for future development is the use of pedigree information when genotype data from parents and/or siblings is also available, which could bolster the accuracy of LD-PGTA at very low coverages. We also note that while not yet tested for these applications, the LD-PGTA framework should generalize to other low-coverage sequencing scenarios, such as prenatal, postnatal, and miscarriage testing.

The genomic resolution of LD-PGTA is a function of multiple factors, including technical variables such as depth and evenness of sequencing coverage, read lengths, and desired confidence level, as well as biological variables such as the magnitudes of local LD and genetic variation. The low genetic diversity of human populations (π ≈ 0.001) is particularly limiting, even with increased sequencing coverage. In practice, an average coverage of 0.01x offers resolution of ~5 Mbp, which corresponds to 10 non-overlapping genomic bins on the shortest human chromosome. At higher depths of coverage (~0.5x), our method permits the mapping of points of transition between different ploidy configurations. Depending on the particular hypotheses under consideration, such transitions could reflect evidence of segmental aneuploidies or—in the case of BPH and SPH trisomy—allow the mapping of meiotic crossover breakpoints on trisomic chromosomes. Beyond the clinical applications previously discussed, this novel output of our method will facilitate future research into the factors influencing meiotic recombination, as well as its impacts on aneuploidy formation—topics of longstanding interest in basic reproductive biology [58].

Standard sequencing-based implementations of PGT-A infer the copy number of each chromosome based on variation in the local depth of coverage, and typically support coverages as low as 0.005x. In contrast, LD-PGTA requires mean coverage of approximately 0.05x to yield high accuracy for any individual embryo, as demonstrated by simulations across a wide range of coverage, read length, and ancestry and admixture scenarios. Nevertheless, when applied to large datasets such as that investigated in our study, global patterns emerge at even lower coverage that offer rough stratification of meiotic and mitotic errors and provide insight into the biological origins of aneuploidy, beyond the binary classification of aneuploid and euploid. As costs of sequencing continue to plummet, application of the method to higher coverage datasets and additional stages of development will further unlock its potential for both research and diagnostic aims.

We envision our method complementing rather than supplanting current coverage-based approaches to PGT-A, whose performance remains superior for tasks such as classification of monosomies and trisomies of few chromosomes. Meanwhile, the application of LD-PGTA can be used to flag haploidies or triploidies that remain invisible to current coverage-based approaches. Additionally, the subclassification of meiotic and mitotic trisomies may prove valuable as orthogonal evidence to distinguish potentially viable mosaic embryos from those possessing lethal meiotic errors. This knowledge is particularly relevant to the many patients with no euploid-testing embryos available for transfer [59]. Together, our method offers a novel approach for extracting useful hidden information from standard preimplantation and prenatal genetic testing data, toward the goal of improving pregnancy outcomes.

## Methods

### Prioritizing informative reads

Broadly, our method overcomes the sparse nature of low-coverage sequencing data by leveraging LD structure of an ancestry-matched reference panel. Measurements of LD require pairwise and higher order comparisons and may thus grow intractable when applied to large genomic regions. To ensure computational efficiency, we developed a scoring algorithm to prioritize reads based on their potential information content, as determined by measuring haplotype diversity within a reference panel at sites that they overlap. We emphasize that the priority score of a read only depends on variation within the reference panel and not on the alleles that the read possesses. The score of a read is calculated as follows:

1. Based on the reference panel, we list all biallelic SNPs that overlap with the read and their reference and alternative alleles.
2. Using the former list, we enumerate all the possible haplotypes. In a region that contain n biallelic SNPs there are 2^*n*^ possible haplotypes.
3. The frequency of each haplotype is estimated from the reference panel by computing the joint frequency of the alleles that comprise each haplotype.
4. We increment the priority score of a read by one for every haplotype with a frequency between *f*_0_ and 1 – *f*_0_.

An example of scoring a read that overlaps with three SNPs appears in Fig. S8 (where *f*_0_ = 0.05). The scoring metric is based on the principle that reads that overlap with potentially informative SNPs with intermediate allele frequencies should receive high priority, as the inclusion of such sites will increase our ability to discern ploidy hypotheses. In the simplest case, where a read overlaps with only a single SNP, the score of the read would be two when the minor allele frequency (MAF) is at least *f*_0_ and otherwise zero. We note that all observed alleles from the same read are considered as originating from the same underlying molecule. Thus, the score should reflect the number of common haplotypes existing in the population at the chromosomal region that overlaps with the read. For a reference panel on the scale of the 1000 Genomes Project (~2500 individuals), 25% - 45% of common SNPs have a nearest neighbor within 35 bp. Hence, even for short reads, the score metric must account for reads that span multiple SNPs.

### Comparing hypotheses with the likelihood ratio

By virtue of LD, observations of a set of alleles from one read may provide information about the probabilities of allelic states in another read that originated from the same DNA molecule (i.e., chromosome). In contrast, when comparing reads originating from distinct homologous chromosomes, allelic states observed in one read will be uninformative of allelic states observed in the other read. As different ploidy configurations are defined by different counts of identical and distinct homologous chromosomes, sparse genotype observations may provide information about the underlying ploidy status, especially when aggregated over large chromosomal regions. For a set of reads aligned to a defined genomic region, we compare the likelihoods of the observed alleles under four competing hypotheses:

1. A single copy of a chromosome, namely monosomy, which may arise by meiotic mechanisms such as non-disjunction, premature separation of sister chromatids, and reverse segregation or mitotic mechanisms such as mitotic non-disjunction and anaphase lag.
2. Two distinct homologous chromosomes, namely disomy, the outcome of normal meiosis, fertilization, and embryonic mitosis.
3. Two identical homologs with a third distinct homolog, denoted as SPH (single parental homolog). SPH may originate from mitotic error (Fig. 1) or rare meiotic errors without recombination [29].
4. Three distinct homologous chromosomes, denoted as BPH (both parental homologs). BPH observed in any portion of a chromosome is a clear indication of meiotic error (Fig. 1) [29].

Our statistical models are based on the premise that the odds of two reads being drawn from the same haplotype differ under the different scenarios. Specifically, for disomy, the odds are 1: 1, for monosomy, the odds are 1: 0, for BPH trisomy, the odds are 1: 2, and for SPH trisomy, the odds are 5: 4 (Fig. 2). If a pair of reads is drawn from identical homologs, the probability of observing the two alleles is given by the joint frequency of these two alleles (i.e., the frequency of the haplotype that they define) in the reference panel. In contrast, if a pair of reads is drawn from distinct homologs, then the probability of observing the two alleles is simply the product of the frequencies of the two alleles in the reference panel:

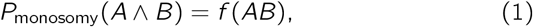

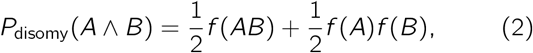

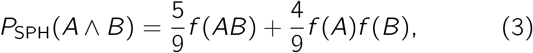

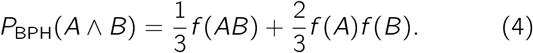

where *f*(*A*) and *f*(*B*) are the frequencies of alleles A and B in the population. Likelihoods of two of the above hypotheses are compared by computing a log-likelihood ratio:

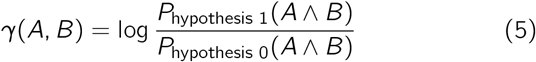

When a read overlaps with multiple SNPs, *f*(*A*) should be interpreted as the joint frequency of all SNP alleles that occur in read *A* (i.e., the frequency of haplotype *A*). Similarly, *f*(*AB*) would denote the joint frequency of all SNP alleles occurring in reads *A* and *B*. The equations above were extended to consider up 18 reads per window, as described in the later section “Generalization to arbitrary ploidy hypotheses”. Estimates of allele and haplotype frequencies from a reference panel do not depend on theoretical assumptions, but rely on the idea that the sample is randomly drawn from a population with similar ancestry. One limitation, which we consider, is that reliable estimates of probabilities near zero or one require large reference panels, such as the 1000 Genome Project [57].

### Quantifying uncertainty by bootstrapping

To quantify uncertainty in our likelihood estimates, we performed m over n bootstrapping by iteratively resampling reads within each window [44]. Resampling was performed without replacement to comply with the assumptions of the statistical models about the odds of drawing two reads from the same haplotype. Thus, in each iteration, only subsets of the available reads can be resampled. Specifically, within each genomic windows, up to 18 reads with a priority score exceeding a defined threshold are randomly sampled with equal probabilities. The likelihood of the observed combination of SNP alleles under each competing hypothesis is then calculated, and the hypotheses are compared by computing a loglikelihood ratio (LLR). The sample mean and the unbiased sample variance (i.e., with Bessel’s correction) of the LLR in each window are calculated by repeating this process using a bootstrapping approach,

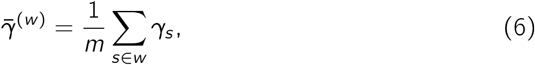

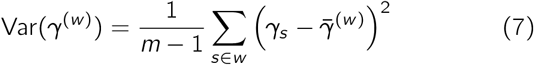

where *γ_s_* is the the log-likelihood ratio for *s*-th subsample of reads from the *w*-th genomic window and *m* is the number of subsamples. Because the number of terms in the statistical models grow exponentially, we subsample at most 18 reads per window. Moreover, accurate estimates of joint frequencies of many alleles requires a very large reference panel. Given the rate of heterozygosity in human populations and the size of the 1000 Genomes dataset, 18 reads is generally sufficient to capture one or more heterozygous SNPs that would inform our comparison of hypotheses.

### Aggregating log likelihood ratios across consecutive windows

Even when sequences are generated according to one of the hypotheses, a fraction of genomic windows will emit alleles that do not support that hypothesis and may even provide modest support for an alternative hypothesis. This phenomenon is largely driven by the sparsity of the data as well as the low rates of heterozygosity in human genomes, which together contribute to random noise. Another possible source of error is a local mismatch between the ancestry of the reference panel and the tested sequence. Moreover, technical errors such as spurious alignment and genotyping could also contribute to poor results within certain genomic regions (e.g., near the centromeres). To overcome this noise, we binned LLRs across consecutive genomic windows, thereby reducing biases and increasing the classification accuracy at the cost of a lower resolution. Specifically, we aggregated the mean LLRs of genomic windows within a larger bin,

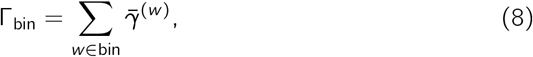

where 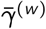 is the mean of the LLRs associated with the *w*-th genomic window. In addition, using the Bienaymé formula, we calculated the variance of the aggregated LLRs,

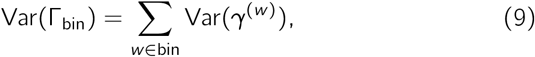

where Var(*γ*^(*w*)^) is the variance of the LLRs associated with a the *w*-th window. For a sufficiently large bin, the confidence interval for the aggregated LLR is 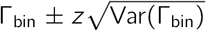, where *z* = Φ^−1^ (1 – *α*) is the *z*-score, Φ is the cumulative distribution function of the standard normal distribution and *C* = 100(1 – 2*α*)% is the confidence level. The confidence level is chosen based on the desired sensitivity vs. specificity. We normalized the aggregated LLRs by the number of genomic windows that comprise each bin, 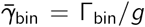. Thus, the variance of the mean LLR per window is 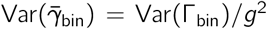. These normalized quantities can be compared across different regions of the genome, as long as the size of the genomic window is the same on average.

### Determining optimal window size

To ensure sufficient data for bootstrap resampling, each genomic window should contain at least one read more than the number of reads in each bootstrap sample. We chose the size of the bootstrap sample as well as the number of bootstrap iterations according to the depth of coverage. Then, we calculated the number of required reads per window and accordingly adjusted the size of the genomic windows.

Pairwise LD in human genomes decays to a quarter of its maximal value over physical distances of 100 kbp, on average [57]. Thus, when (a) the distance between consecutive observed alleles exceed 100 kbp or (b) a genomic window reaches 350 kbp and does not meet the minimal required number of reads, it is dismissed. An adaptive sliding window possesses advantages over a fixed length window in that it (a) accounts for GC-poor and GC-rich regions of a genome, which tend to be sequenced at lower depths of coverage using Illumina platforms [60] and (b) accounts for varying densities of SNPs across the genome [61].

### Simulating meiotic- and mitotic-origin trisomies

To simulate trisomies, we constructed synthetic samples comprised of combinations of three sampled phased haplotypes from the 1000 Genomes Project [57]. These phased haplotypes were extracted from VCF files to effectively form a pool of haploid sequences from which to draw trisomies.

We first assigned full chromosomes or chromosome segments to BPH or SPH trisomy states. We note that true meiotic trisomies exhibit a mixture of BPH and SPH segments, while mitotic trisomies exhibit SPH over their entire length [29]. The BPH and SPH regions of simulated trisomic chromosomes were determined as follows:

1. We assumed that the number of transitions between BPH and SPH regions was equal to the number of meiotic crossover events, on average, per autosome. Thus, for chromosomes 1 – 6, 7 – 12 and 13 – 22 we simulated are 3,2 and 1 transition, respectively.
2. When simulating trisomies of meiosis II origin, we required that the region around the centromere reflect the SPH hypothesis.
3. For simplicity, we assumed homogeneity in the frequency of meiotic crossovers throughout the genome (excluding centromeres), thus drawing transition points (between SPH and BPH) from a uniform distribution.

Reads were simulated by selecting a random position along the chromosome from a uniform distribution, representing the midpoint of an aligned read with a given length. Based on the selected position, one out of the three haplotypes was drawn from a discrete distribution,

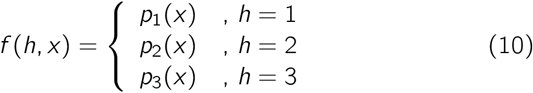

where the probability of haplotype *h* depends on the position of the read, *x*. When the read overlaps with a BPH region, all three haplotypes have the same probability, *p*_1_ = *p*_2_ = *p*_3_. For an SPH region, the first haplotype is twice as likely as the second haplotype, *p*_2_ = 2*p*_1_, while the third haplotype is absent, *p*_3_ = 0.

From the selected haplotype, *h*, a segment of length *l* that is centered at the selected chromosomal position, *x*, is added to simulated data, mimicking the process of short-read sequencing. This process of generating simulated sequencing data is repeated until the desired depth of coverage is attained.

### Evaluating model performance on simulated data

We developed a classification scheme to infer the ploidy status of each genomic bin. Each class is associated with a hypothesis about the number of distinct homologs and their degeneracy (i.e., copy number of non-unique homologs). To this end, we compare pairs of hypotheses by computing log likelihood ratios (LLRs) of competing statistical models. In general, the specific models that we compare are informed by orthogonal evidence obtained using standard coverage-based approaches to aneuploidy detection (i.e., tag counting).

The confidence interval for the mean LLR is 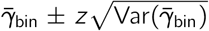, and *z* is referred to as the *z*-score. Thus, we classify a bin as exhibiting support for hypothesis 1 when

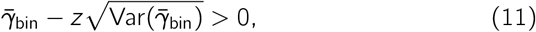

and for hypothesis 0 when

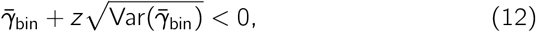

where the first (second) criterion is equivalent to requiring that the bounds of the confidence interval lie on the positive (negative) side of the number line.

For a given depth of coverage and read length, we simulate an equal number of sequences generated according to hypothesis 0 and 1, as explained in the previous section. Based on these simulations, we generate two receiver operating characteristic (ROC) curves for each bin. The first ROC curve is produced by defining true positives as simulations where sequences generated under hypothesis 0 are correctly classified. For the second ROC curve, true positives are defined as simulations where sequences generated under hypothesis 1 are correctly classified. These two ROC curves can be combined into a single curve, which we term “balanced ROC curve”. The balanced true and false positive rates for a bin are defined as

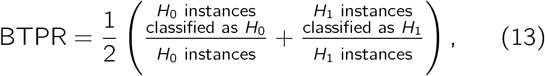

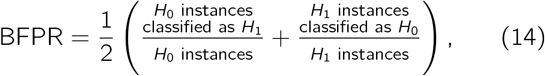

where a H_i_ instance with *i* =1,2 is a sequence that was generated under hypothesis *i*.

The balanced ROC curve is particularly suited for classification tasks with three possible classes: *H*_0_, *H*_1_ and ambiguous. The ambiguous class contains all instances that do not fulfill the criteria in Eqs. (11) and (12), i.e., instances where the boundaries of the confidence interval span zero. This classification scheme allows us to optimize the classification of both *H*_0_ and *H*_1_ instances, at the expense of leaving ambiguous instances. The advantage of this optimization is a reduction in the rate of spurious classification. To generate each curve, we varied the confidence level and the number of bins.

### Generalization to arbitrary ploidy hypotheses

While the aforementioned ploidy hypotheses (monosomy, disomy, SPH trisomy, and BPH trisomy) are most relevant to our study, our method can be generalized to arbitrary ploidy hypotheses and an arbitrary number of observed alleles. Consider *m* reads from an autosome with a copy number of *n*, containing *l* unique homologs. We list all the possible ways in which reads may emanate from the different homologs. For example, in the case of two reads, denoted by *A* and *B*, and two unique homologs, the list contains four configurations, i.e., ({*A*}, {*B*}), ({*B*}, {*A*}), ({*AB*}, 0), (0, {*AB*}). We assign a weight to each configuration in the list. The weight depends on the degeneracy of each homolog. For example, the presence of two identical homologs means that the degeneracy of this homolog is two.

For a each configuration, we list the number of reads that were assigned to each homolog along with the degeneracy of these homologs. Moreover, the *i*^th^ element in the list is (*r_i_, d_i_*) where *r_i_* is the number of reads assigned to the *i*^th^ homolog and *d_i_* is the degeneracy of this homolog. We also note that for each configuration, the relations 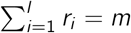 and 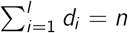 hold. Then, the weight associated with a configuration is 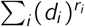.

All configurations that share the same partitioning of reads, regardless of the homolog to which the subset was assigned, are grouped together. For example, in the case of two reads and two unique homologs there are two possible partitions, i.e., {{*A*}, {*B*}} and {*AB*}, 0}. We associate each partition of reads with the total weight of all the configurations that share the same partition. In the case of two reads from a chromosome with two unique homologs out of three, we have (*AB*, 5) and (*A|B*, 4).

Each partition together with its associated weight, contributed a term to the statistical model. The term is a product of a normalized weight and joint frequencies. The normalization factor is *n^−m^* and each joint frequency corresponds to a subset of reads, e.g., using the partitions list from the former example we obtain 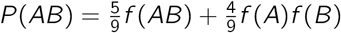.

### Efficient encoding of statistical models

The number of terms in the statistical models grows exponentially with the number of alleles, and it is thus necessary to write the models in their simplest forms and encode them efficiently.

In order to identify common multiples in the statistical model, we group partitions of reads according to their subset with the smallest cardinality. For example, the partitions {{*A*}, {*B,C*}, {*D, E*}} and {{*A*}, {*B, E*}, {*C, D*}} share the subset {*A*}, which correspond to the frequency *f*(*A*).

The partitioning of reads can be encoded efficiently using the occupation basis. In this representation, all reads are enumerated. Each subset of reads is represented by a binary sequence, where the *i*^th^ element is one when the *i*^th^ read is included in the subset and zero otherwise. In other words, bits in a binary sequence denote whether a read is included in the subset. For example, when the first two reads are associated with the same homolog and the third read is associated with another homolog then (1,2), (3) corresponds to (0,1,1), (1,0, 0).

### Modeling admixture in the previous generation

Admixed individuals constitute a considerable portion of contemporary societies, and indeed all genomes possess admixture at varying scales and degrees. Here we modify the statistical models to account for admixture between two defined populations. A chromosome of an individual of recently admixed ancestry resembles a mosaic of chromosomal segments, each derived from a particular ancestral population [62]. Thus, the associated statistical models should account for the possibility of admixed ancestry in the selection of appropriate reference panels.We first consider an individual descended from parents of distinct ancestries (hereafter termed “recent admixed”). In the case of monosomy, the observed homolog is equally likely to have originated from either of the two parental populations,

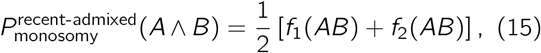

where *f*_1_ and *f*_2_ are joint frequency distributions that are associated with each respective population. For disomy, each homolog is inherited from a parent deriving from a different population,

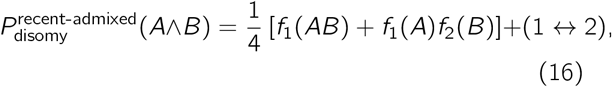

where (1 ↔ 2) is equal to the sum of all the other terms in the expression with the indices 1 and 2 exchanged, e.g., *f*_1_(*A*)*f*_2_(*B*) + (1 ↔ 2) = *f*_1_(*A*)*f*_2_(*B*) + *f*_2_(*A*)*f*_1_(*B*). Considering SPH trisomy, the third non-unique homolog originated from either of the two ancestral populations with equal probability and thus,

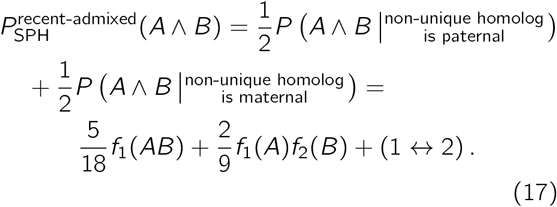

Similarly, for the BPH trisomy hypothesis we have

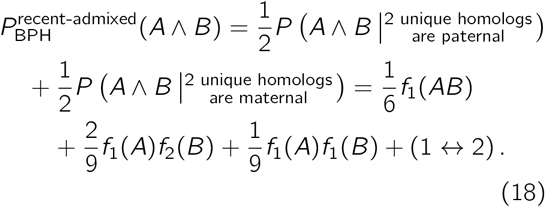

### Extending the recent admixture model to an arbitrary number of reads

The monosomy, disomy, and SPH trisomy statistical models for recent admixed individuals and an arbitrary number of reads can be obtained from the corresponding statistical models for non-admixed individuals. We start with a statistical model for a non-admixed individual, where the SPH model for 3 reads is

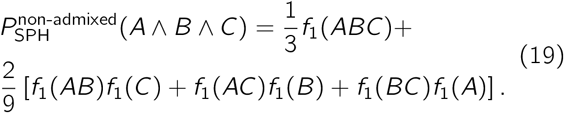

We re-express each term with arbitrary frequency distributions, i.e., *f*_1_ → *f_i_*, *f*_1_*f*_1_ → *f_i_f_j_*. Then, we multiply the expression by (1 – *δ_ij_*)/2 and sum over *i* and *j*. Here and in what follows, *δ_ij_* and *δ_i,j,k_* are the Kronecker delta and a generalization, respectively, and are defined as

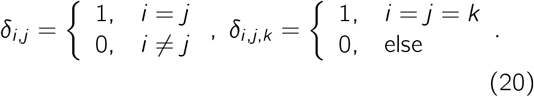

Following these three steps, we obtain the recent admixed SPH model for 3 reads,

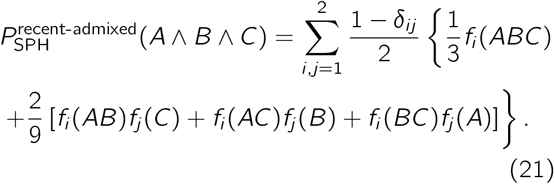

Statistical models of BPH trisomy for recent admixed individuals and an arbitrary number of reads can also be obtained from the corresponding statistical models for non-admixed individuals. We start with a statistical model for a non-admixed individual, where the BPH model for 3 reads is

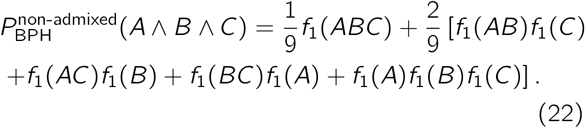

We continue by re-expressing each term with arbitrary frequency distributions, i.e., *f*_1_ → *f_i_, f*_1_*f*_1_ → *f_i_f_j_* and *f*_1_*f*_1_*f*_1_ → *f_i_f_j_f_k_*. Then, we multiply the expression by (1 – *δ_i,j,k_*)/6 and sum over *i,j* and *k*. In keeping with our previous example we obtain

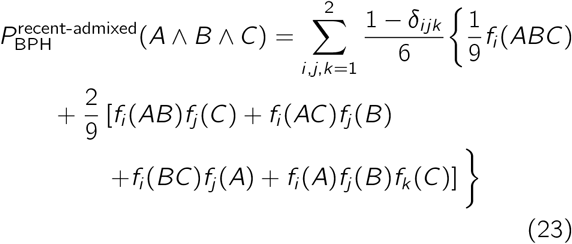

### Generalized models for more distant and arbitrary admixture scenarios

While the recent admixture models consider the specific case of admixture involving parents from distinct populations, admixture could also occur in earlier generations and/or involve multiple populations, the details of which may be unknown. It is therefore important to also consider amendments to the models to account for such arbitrary admixture scenarios, where only the overall ancestry proportions may be known, for example as estimated by popular methods [63–65].

The statistical models for these arbitrary admixture scenarios are based on the assumption that each distinct parental haplotype is drawn from an ancestral population with a probability equal to the ancestry proportion of the tested individual that is associated with that population. Thus, the probability of observing alleles that are associated with a single parental haplotype (i.e., monosomy) is a linear combination of their joint-frequencies in various ancestral populations,

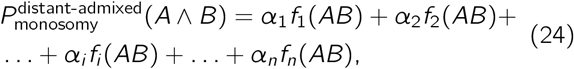

where *α_i_* and *f_i_*(*AB*) are the admixture proportion and the joint-frequency of the alleles *A* and *B* with respect to the *i*-th ancestral population and 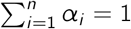.

All such statistical models can then be extended to an arbitrary number of reads by adapting the corresponding statistical models for non-admixed individuals. This is achieved by replacing each allele frequency distribution, *f* by the combination *α*_1_*f*_1_ + *α*_2_*f*_2_ + … + *α_i_f_i_* + … + *α_n_f_n_*. Here *α_i_* is the probability that the alleles originated from the *i*-th population, *f_i_* is the allele frequency distribution for the *i*-th population and 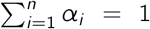. For example, the BPH model for admixture that involves two ancestral populations can be expressed as 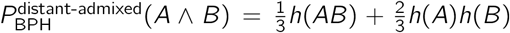 where *h*(*X*) ≡ *α*_1_*f*_1_(*X*) + (1 – *α*_1_)*f*_2_(*X*). Admixture models for BPH, SPH, disomy, and monosomy for up to 4 reads are explicitly provided in the Supplementary Materials. In addition, balanced ROC curves that demonstrate the performance of the classifier on simulated trisomy of Chromosome 21 can also be found in the Fig. S9.

### Accounting for the rate of heterozygosity

One important feature of likelihood ratios is that the ratio rather than individual likelihoods should be the focus of interpretation. When a read overlaps with a common SNP and the embryo is homozygous at this locus, the read is uninformative for distinguishing trisomy hypotheses. We address this limitation in several ways:

1. The number of reads required to distinguish trisomy hypotheses is largely determined by the rate of heterozygosity in human genomes. Informative (heterozygous) sites in the target sample are unknown *a priori*. Even sites of common variation in the reference panel will frequently be homozygous in the target sample, necessitating sampling of many reads to capture informative sites.
2. The probability of observing genetic differences between haplotypes increases with the length of the region investigated. All alleles observed within a single read necessarily originate from the same molecule. Our model accounts for the possibility that a read overlaps multiple SNPs, any one of which may be informative for distinguishing haplotypes. Longer reads thus provide direct readout of haplotypes, improving performance of our method.
3. We use a scoring algorithm to prioritize reads that overlap with diverse haplotypes within the a reference panel, as described in the scoring section of the Methods. It is worth emphasizing that at extremely low-coverages, rare alleles play an important role in distinguishing trisomy hypotheses. Thus, one should avoid narrowing the bandwidth, Δ*f* = 1 – 2*f*_0_ of the scoring metric beyond what is necessary to ensure computational efficiency.
4. Because a window might not contain reads that overlap with heterozygous sites, we use binning to aggregate the mean LLR across multiple consecutive windows. This procedure, which reduces biases inherent to low-coverage sequence data, especially in species and populations with low heterozygosity, is described in the aggregation section of the Methods.

For both the BPH and SPH statistical models, we checked whether small changes in the degeneracies of the unique homologs can compensate for low rates of heterozygosity in triploids. For example, instead of three unique homologs with equal degeneracies, (1, 1, 1) one can consider the case (1.4, 1.2, 1), which is equivalent to (7, 6, 5). We observed that for statistical models of at least three reads, small changes of the degeneracies did not substantially affect the percentage windows that were correctly classified. This underscores the robustness of the statistical models and suggests that the number of unique homologs is the most important parameter defining each hypothesis. One implication is that our approach is less efficient at distinguishing SPH trisomy from disomy, as both scenarios involve two unique homologs (albeit in different proportions). We therefore emphasize that LD-PGTA is best considered as a complement to, rather than a replacement for, standard coverage-based methods.

### Implications of overlapping reads

Even at extremely low coverage, the probability of two reads to overlap is not negligible. Although overlaps of two reads are not sufficient to distinguish between ploidy hypotheses, they reduce the complexity of the calculations:

1. When two different alleles are observed at the same SNP, it means that they necessarily originated from different haplotypes. Thus, eliminating some of the terms in the statistical models.
2. When the same allele is observed twice, it reduces the dimensionality of the joint frequency, e.g., *f*(*AA*) = *f*(*A*).

### Mapping meiotic crossovers

Transitions between the BPH and SPH hypotheses along the chromosome indicate the locations of meiotic crossovers. We approximated each crossover location as the center between two adjacent bins, where one is classified as BPH and the second as SPH. This approximation becomes more precise as the size of the bins is reduced and the confidence interval becomes smaller.

### Detecting haploidy and triploidy

A key aim of our method is the reclassification of sequences that were initially identified as euploid by standard coverage based approaches as exhibiting potential abnormalities of genome-wide ploidy. Specifically, coverage-based methods are based on the correlation between the depth of coverage of reads aligned to a genomic region and the copy number of that region. In the cases of (near-) haploidy and triploidy, no such correlation is evident because (nearly) all chromosomes exhibit the same abnormality, resulting in erroneous classification.

To this end, following the binning procedure that was introduced in Eqs. (8) and (9), we aggregate the LLRs of each genomic window along the entire genome. In order to identify haploids, we test for cases where the confidence interval lies on the positive side of the number line, as formulated in Eq. (11).

In the idealized case where triploids are composed of entirely BPH sequence, triploids are easily distinguished from diploids. However, true cases of triploidy generally originate from retention of the polar body, leading to a more realistic scenario in which regions of BPH are relegated to the ends of chromosomes, while the rest of the genome exhibits SPH. The remedy is to introduce a binary classifier that assesses whether there is enough evidence of genome-wide BPH to question the hypothesis of genome-wide diploidy,

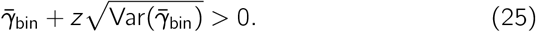

In other words, it is sufficient to show that the confidence interval spans the zero in order to classify the case as a triploid. We evaluated this criterion by applying our method to simulations of triploid embryos with realistic tracts of BPH and SPH trisomy. ROC curves constructed from these simulations demonstrate near-ideal performance at coverage ≥0.05x with a correctly specified reference panel (Figs. S8a and S8b).

## Acknowledgements

Thank you to the Origins of Aneuploidy Research Consortium, Shai Carmi, Alexander Zaranek, as well as members of the McCoy lab for constructive feedback. Thanks also to staff of the Maryland Advanced Research Computing Center for computing support.

## Funding

Research reported in this publication was supported by National Institute of General Medical Sciences of the National Institutes of Health under award number R35GM133747. The content is solely the responsibility of the authors and does not necessarily represent the official views of the National Institutes of Health.

## Availability of data and materials

Software for implementing our method and reproducing all analysis is available at: https://github.com/mccoy-lab/LD-PGTA. Bluefuse Multi aneuploidy data and LD-PGTA results for all analyzed chromosomes are available in the same repository.

## Ethics approval and consent to participate

The Homewood Institutional Review Board of Johns Hopkins University determined that this work does not qualify as federally-regulated human subjects research (HIRB00011431).

## Competing interests

D.A., M.V., and R.C.M. are co-inventors of the method described herein, which is the subject of a provisional patent application by Johns Hopkins University.

## Consent for publication

All authors have read and approved the manuscript.

## Authors’ contributions

D.A.: Methodology, Software, Formal analysis, Investigation, Data curation, Writing - Original Draft, Writing - Review & Editing, Visualization; S.M.Y.: Writing - Original Draft, Writing - Review & Editing; A.R.V.: Data collection, Data curation; F.L.B.: Data collection, Data curation. C.G.Z.: Data collection, Data curation; M.V.: Conceptualization, Data collection, Data curation, Writing - Review & Editing; R.C.M.: Conceptualization, Writing - Original Draft, Writing - Review & Editing, Visualization, Supervision, Funding acquisition

## Notes

https://github.com/mccoy-lab/LD-PGTA

### Summary of Updates

Updated notation for admixture models to avoid unintended connotation of an experimental cross. Corrected formatting of several references.

