## Supplementary Materials for "Haplotype-aware inference of human chromosome abnormalities"

#### Statistical models for non-admixed ancestry

The monosomy, disomy, SPH and BPH statistical models for individuals with non-admixed ancestry for between two to four reads are given below:

##### *Two Reads*

$$P^{\text{non-admixed}}_{\text{monosomy}}(A \wedge B) = f(AB)$$

$$P^{\text{non-admixed}}_{\text{disomy}}(A \wedge B) = \frac{1}{2}\left[f(AB) + f(A)f(B)\right]$$

$$P^{\text{non-admixed}}_{\text{SPH}}(A \wedge B) = \frac{5}{9}f(AB) + \frac{4}{9}f(A)f(B)$$

$$P^{\text{non-admixed}}_{\text{BPH}}(A \wedge B) = \frac{1}{3}f(AB) + \frac{2}{3}f(A)f(B)$$

##### *Three reads*

$$P^{\text{non-admixed}}_{\text{monosomy}}(A \wedge B \wedge C) = f(ABC)$$

$$P^{\text{non-admixed}}_{\text{disomy}}(A \wedge B \wedge C) = \frac{1}{4}\left[f(ABC) + f(AB)f(C) + f(AC)f(B) + f(BC)f(A)\right]$$

$$P^{\text{non-admixed}}_{\text{SPH}}(A \wedge B \wedge C) = \frac{1}{3}f(ABC) + \frac{2}{9}\left[f(AB)f(C) + f(AC)f(B) + f(BC)f(A)\right]$$

$$P^{\text{non-admixed}}_{\text{BPH}}(A \wedge B \wedge C) = \frac{1}{9}f(ABC) + \frac{2}{9}\left[f(AB)f(C) + f(AC)f(B) + f(BC)f(A) + f(A)f(B)f(C)\right]$$

##### *Four reads*

$$P^{\text{non-admixed}}_{\text{monosomy}}(A \wedge B \wedge C \wedge D) = f(ABCD)$$

$$\begin{aligned}P^{\text{non-admixed}}_{\text{disomy}}(A \wedge B \wedge C \wedge D) =& \frac{1}{8}\big[f(ABCD) + f(ABC)f(D) + f(BCD)f(A) + f(ACD)f(B) + \\ & f(ABD)f(C) + f(AB)f(CD) + f(AC)f(BD) + f(AD)f(BC)\big]\end{aligned}$$

$$\begin{aligned}P^{\text{non-admixed}}_{\text{SPH}}(A \wedge B \wedge C \wedge D) =& \frac{17}{81}f(ABCD) + \frac{10}{81}\big[f(ABC)f(D) + f(BCD)f(A) + f(ACD)f(B) + \\ & f(ABD)f(C)) + \frac{8}{81}(f(AB)f(CD) + f(AD)f(BC) + f(AC)f(BD)\big]\end{aligned}$$

$$P^{\text{non-admixed}}_{\text{BPH}}(A \wedge B \wedge C \wedge D) = \frac{1}{27} f(ABCD) + \frac{2}{27}\big[f(AB)f(C)f(D) + f(A)f(BD)f(C) + f(A)f(BC)f(D) + f(AC)f(B)f(D) + f(A)f(B)f(CD) + f(AD)f(B)f(C) + f(ABC)f(D) + f(A)f(BCD) + f(ACD)f(B) + f(ABD)f(C) + f(AB)f(CD) + f(AD)(BC) + f(AC)f(BD)\big]$$

The function $f$ is the joint frequency distributions, which is derived from a reference panel. The alleles $A, B, C$ and $D$ are represented by vectors of chromosomal position and base pair. For brevity we use the notation $f(ABC\ldots Z) \equiv f(A, B, C, \ldots, Z)$, i.e. when $AB$ appears as an argument of a function it reflects two distinct arguments, $A$ and $B$ and not a scalar product of the two.

### Statistical models for admixture in the previous generation

The monosomy, disomy, SPH and BPH statistical models for recent admixed individuals for between two to four reads are given below:

#### *Two Reads*

$$P^{\text{recent-admixed}}_{\text{monosomy}}(A \wedge B) = \frac{1}{2}\big[f_1(AB) + f_2(AB)\big]$$

$$P^{\text{recent-admixed}}_{\text{disomy}}(A \wedge B) = \frac{1}{4}\big[f_1(AB) + f_1(A)f_2(B) + f_2(A)f_1(B) + f_2(AB)\big]$$

$$P^{\text{recent-admixed}}_{\text{SPH}}(A \wedge B) = \frac{2}{9}\big[f_1(A)f_2(B) + f_2(A)f_1(B)\big] + \frac{5}{18}\big[f_1(AB) + f_2(AB)\big]$$

$$P^{\text{recent-admixed}}_{\text{BPH}}(A \wedge B) = \frac{1}{9}\big[f_1(B)f_1(A) + f_2(B)f_2(A)\big] + \frac{1}{6}\big[f_1(AB) + f_2(AB)\big] + \frac{2}{9}\big[f_1(B)f_2(A) + f_2(B)f_1(A)\big]$$

#### *Three reads*

$$P^{\text{recent-admixed}}_{\text{monosomy}}(A \wedge B \wedge C) = \frac{1}{2}\big[f_1(ABC) + f_2(ABC)\big]$$

$$P^{\text{recent-admixed}}_{\text{disomy}}(A \wedge B \wedge C) = \frac{1}{8}\big[f_1(A)f_2(BC) + f_1(B)f_2(AC) + f_1(C)f_2(AB) + f_1(ABC)\big] + (1 \leftrightarrow 2)$$

$$P^{\text{recent-admixed}}_{\text{SPH}}(A \wedge B \wedge C) = \frac{1}{9}\big[f_1(A)f_2(BC) + f_1(B)f_2(AC) + f_1(C)f_2(AB)\big] + \frac{1}{6} f_1(ABC) + (1 \leftrightarrow 2)$$

$$P^{\text{recent-admixed}}_{\text{BPH}}(A \wedge B \wedge C) = \frac{1}{27}\big[f_1(A)f_2(B)f_2(C) + f_1(A)f_1(B)f_2(C) + f_1(A)f_2(B)f_1(C) + f_1(A)f_1(BC) + f_1(B)f_1(AC) + f_1(C)f_1(AB)\big] + \frac{2}{27}\big[f_1(A)f_2(BC) + f_1(B)f_2(AC) + f_1(C)f_2(AB)\big] + \frac{1}{18} f_1(ABC) + (1 \leftrightarrow 2)$$

*Four reads*

$$P_{\text{monosomy}}^{\text{recent-admixed}}(A \wedge B \wedge C) = \frac{1}{2}\left[f_1(ABCD) + f_2(ABCD)\right]$$

$$\begin{aligned} P_{\text{disomy}}^{\text{recent-admixed}}(A \wedge B \wedge C) =& \frac{1}{16}\left[f_1(A)f_2(BCD) + f_1(B)f_2(ACD) + f_1(C)f_2(ABD) + f_1(D)f_2(ABC) + \right. \\ & \left. f_1(AB)f_2(CD) + f_1(AC)f_2(BD) + f_1(AD)f_2(BC) + f_1(ABCD)\right] + (1 \leftrightarrow 2) \end{aligned}$$

$$\begin{aligned} P_{\text{SPH}}^{\text{recent-admixed}}(A \wedge B \wedge C) =& \frac{4}{81}\left[f_1(AB)f_2(CD) + f_1(AC)f_2(BD) + f_1(AD)f_2(BC)\right] + \frac{5}{81}\left[f_1(A)f_2(BCD) + \right. \\ & \left. f_1(B)f_2(ACD) + f_1(C)f_2(ABD) + f_1(D)f_2(ABC)\right] + \frac{17}{162}f_1(ABCD) + (1 \leftrightarrow 2) \end{aligned}$$

$$\begin{aligned} P_{\text{BPH}}^{\text{recent-admixed}}(A \wedge B \wedge C \wedge D) =& \frac{1}{81}\Big\{f_1(A)\big[f_1(BCD) + f_1(B)f_2(CD) + f_2(B)f_1(CD) + f_2(B)f_2(CD) + \\ & f_1(C)f_2(BD) + f_2(C)f_1(BD) + f_2(C)f_2(BD) + f_1(D)f_2(BC) + f_2(D)f_1(BC) + \\ & f_2(D)f_2(BC)\big] + f_1(B)\big[f_1(ACD) + f_1(C)f_2(AD) + f_2(C)f_1(AD) + f_2(C)f_2(AD) + \\ & f_1(D)f_2(AC) + f_2(D)f_1(AC) + f_2(D)f_2(AC)\big] + f_1(C)\big[f_1(ABD) + f_1(D)f_2(AB) + \\ & f_2(D)f_1(AB) + f_2(D)f_2(AB)\big] + f_1(D)f_1(ABC) + f_1(AB)f_1(CD) + \\ & f_1(AC)f_1(BD) + f_1(AD)f_1(BC)\Big\} + \frac{2}{81}\big[f_1(A)f_2(BCD) + f_1(B)f_2(ACD) + \\ & f_1(C)f_2(ABD) + f_1(D)f_2(ABC) + f_1(AB)f_2(CD) + f_1(AC)f_2(BD) + \\ & f_1(AD)f_2(BC)\big] + \frac{1}{54}f_1(ABCD) + (1 \leftrightarrow 2) \end{aligned}$$

The notation $(1 \leftrightarrow 2)$ is used to represent the sum of all the other terms in the expression with the indices 1 and 2 exchanged, e.g., $f_1(A)f_2(B) + (1 \leftrightarrow 2) = f_1(A)f_2(B) + f_2(A)f_1(B)$.

### Statistical models for more distant and arbitrary admixture scenarios

The monosomy, disomy, SPH and BPH statistical models for individuals with distant and arbitrary admixed ancestry for between two to four reads are given below:

*Two Reads*

$$P_{\text{monosomy}}^{\text{distant-admixed}}(A \wedge B) = \sum_{i=1}^{2} \alpha_i f_i(AB)$$

$$P_{\text{disomy}}^{\text{distant-admixed}}(A \wedge B) = \sum_{i,j=1}^{2} \frac{\alpha_i \alpha_j}{4}\left[f_i(AB) + f_j(AB) + f_i(A)f_j(B) + f_i(B)f_j(A)\right]$$

$$P_{\text{SPH}}^{\text{distant-admixed}}(A \wedge B) = \sum_{i,j=1}^{2} \frac{\alpha_i \alpha_j}{9}\left\{4f_i(AB) + f_j(AB) + 2\left[f_i(A)f_j(B) + f_i(B)f_j(A)\right]\right\}$$

$$P_{\text{BPH}}^{\text{distant-admixed}}(A \wedge B) = \sum_{i,j,k=1}^{2} \frac{\alpha_i \alpha_j \alpha_k}{9}\left\{f_i(AB) + f_j(AB) + f_k(AB) + 2\left[f_i(A)f_j(B) + f_j(A)f_k(B) + f_k(A)f_i(B)\right]\right\}$$

*Three reads*

$$P^{\text{distant-admixed}}_{\text{monosomy}}(A \wedge B \wedge C) = \sum_{i=1}^{2} \alpha_i f_i(ABC)$$

$$P^{\text{distant-admixed}}_{\text{disomy}}(A \wedge B \wedge C) = \sum_{i,j=1}^{2} \frac{\alpha_i \alpha_j}{8} \{f_i(ABC) + f_j(ABC) + f_i(AB)f_j(C) + f_i(AC)f_j(B) + f_i(BC)f_j(A) + \\ + f_i(A)f_j(BC) + f_i(B)f_j(AC) + f_j(AB)f_i(C)\}$$

$$P^{\text{distant-admixed}}_{\text{SPH}}(A \wedge B \wedge C) = \sum_{i,j=1}^{2} \frac{\alpha_i \alpha_j}{27} \{8f_i(ABC) + f_j(ABC) + 4\left[f_i(AB)f_j(C) + f_i(AC)f_j(B) + f_i(BC)f_j(A)\right] + \\ +2\left[f_i(A)f_j(BC) + f_i(B)f_j(AC) + f_i(C)f_j(AB)\right]\}$$

$$P^{\text{distant-admixed}}_{\text{BPH}}(A \wedge B \wedge C) = \sum_{i,j,k=1}^{2} \frac{\alpha_i \alpha_j \alpha_k}{27} \{f_i(ABC) + f_j(ABC) + f_k(ABC) + 2\left[f_i(A)f_j(B)f_k(C) + \right. \\ f_k(A)f_i(B)f_j(C) + f_j(A)f_k(B)f_i(C) + f_i(AB)f_j(C) + f_j(AB)f_k(C) + \\ f_k(AB)f_i(C) + f_i(AC)f_j(B) + f_j(AC)f_k(B) + f_k(AC)f_i(B) + f_i(BC)f_j(A) + \\ \left. f_j(BC)f_k(A) + f_k(BC)f_i(A)\right]\}$$

*Four reads*

$$P^{\text{distant-admixed}}_{\text{monosomy}}(A \wedge B \wedge C \wedge D) = \sum_{i=1}^{2} \alpha_i f_i(ABCD)$$

$$P^{\text{distant-admixed}}_{\text{disomy}}(A \wedge B \wedge C) = \sum_{i,j,k=1}^{2} \frac{\alpha_i \alpha_j}{16} \left[f_i(ABCD) + f_j(ABCD) + f_i(ABC)f_j(D) + f_i(ABD)f_j(C) + f_i(ACD)f_j(B) \right. \\ f_i(BCD)f_j(A) + f_i(AB)f_j(CD) + f_i(AC)f_j(BD) + f_i(AD)f_j(BC) + f_i(BC)f_j(AD) + \\ f_i(BD)f_j(AC) + f_i(CD)f_j(AB) + f_i(A)f_j(BCD) + f_i(B)f_j(ACD) + \\ \left. f_i(C)f_j(ABD) + f_i(D)f_j(ABC)\right]$$

$$P^{\text{distant-admixed}}_{\text{SPH}}(A \wedge B \wedge C) = \sum_{i,j,k=1}^{2} \frac{\alpha_i \alpha_j}{81} \{16f_i(ABCD) + f_j(ABCD) + 8\left[f_i(ABC)f_j(D) + f_i(ABD)f_j(C) + \right. \\ \left. f_i(ACD)f_j(C) + f_i(BCD)f_j(A)\right] + 4\left[f_i(AB)f_j(CD) + f_i(AC)f_j(BD) + f_i(AD)f_j(BC) + \right. \\ \left. f_i(BC)f_j(AD) + f_i(BD)f_j(AC) + f_i(CD)f_j(AB)\right] + 2\left[f_i(A)f_j(BCD) + \right. \\ \left. f_i(B)f_j(ACD) + f_i(C)f_j(ABD) + f_i(D)f_j(ABC)\right]\}$$

$$
\begin{aligned}
P_{\text{BPH}}^{\text{distant-admixed}}(A \wedge B \wedge C \wedge C \wedge D) = \quad & \sum_{i,j,k=1}^{2} \frac{\alpha_i \alpha_j \alpha_k}{81} \{ f_i(ABCD) + f_j(ABCD) + f_k(ABCD) \\
& +2\,[f_i(AB)f_j(C)f_k(D) + f_k(AB)f_i(C)f_j(D) + f_j(AB)f_k(C)f_i(D) + \\
& f_i(AC)f_j(B)f_k(D) + f_k(AC)f_i(B)f_j(D) + f_j(AC)f_k(B)f_i(D) + \\
& f_i(A)f_j(BC)f_k(D) + f_k(A)f_i(BC)f_j(D) + f_j(A)f_k(BC)f_i(D) + \\
& f_i(A)f_j(BD)f_k(C) + f_k(A)f_i(BD)f_j(C) + f_j(A)f_k(BD)f_i(C) + \\
& f_i(A)f_j(B)f_k(CD) + f_k(A)f_i(B)f_j(CD) + f_j(A)f_k(B)f_i(CD) + \\
& f_i(AD)f_j(B)f_k(C) + f_k(AD)f_i(B)f_j(C) + f_j(AD)f_k(B)f_i(C) + \\
& f_i(ABC)f_j(D) + f_j(ABC)f_k(D) + f_k(ABC)f_i(D) + \\
& f_i(ABD)f_j(C) + f_j(ABD)f_k(C) + f_k(ABD)f_i(C) + \\
& f_i(AB)f_j(CD) + f_j(AB)f_k(CD) + f_k(AB)f_i(CD) + \\
& f_i(ACD)f_j(B) + f_j(ACD)f_k(B) + f_k(ACD)f_i(B) + \\
& f_i(A)f_j(BCD) + f_j(A)f_k(BCD) + f_k(A)f_i(BCD) + \\
& f_i(AC)f_j(BD) + f_j(AC)f_k(BD) + f_k(AC)f_i(BD) + \\
& f_i(AD)f_j(BC) + f_j(AD)f_k(BC) + f_k(AD)f_i(BC)] \}
\end{aligned}
$$

### Supplemental Tables

Table S1 (Top) Table of weights for 3 reads under the SPH scenario. (Bottom) Based on the table a statistical model was derived.

| Case Description | Haploid 1 (no degeneracy) | Haploid 2 (degeneracy of 2) | Weight | Partition | Total Weight |
|---|---|---|---|---|---|
| Only last two reads matches the same haplotype | $A$<br>$BC$ | $BC$<br>$A$ | $1^1 \times 2^2 = 4$<br>$1^2 \times 2^1 = 2$ | $A\|BC$ | 6 |
| Only first and third reads matches the same haplotype | $B$<br>$AC$ | $AC$<br>$B$ | 4<br>2 | $B\|AC$ | 6 |
| Only first two reads matches the same haplotype | $C$<br>$AB$ | $AB$<br>$C$ | 4<br>2 | $C\|AB$ | 6 |
| All three reads matches the same haplotype | $-$<br>$ABC$ | $ABC$<br>$-$ | $1^0 \times 2^3 = 8$<br>$1^3 \times 2^0 = 1$ | $ABC$ | 9 |
| | | | | | 27 |

$$P_{\text{SPH}}(ABC) = \tfrac{6}{27}\big(f(A)f(BC) + f(B)f(AC) + f(C)f(AB)\big) + \tfrac{9}{27}f(ABC)$$

Table S2 Statistics of the simulated non-admixed triploids, which were used for generating the balanced ROC curve in Fig. 3c. The table includes information about bin sizes, GW sizes, number of GWs per bin and number or reads per GW. All simulated sequences consist of reads 36 bp in length.

| Depth | Ancestry | Reference Panel | Bin size (kbp) | | GW size (kbp) | | GW per Bin | | Reads per GW | | Sampled Reads |
|---|---|---|---|---|---|---|---|---|---|---|---|
| | | | Mean | SD | Mean | SD | Mean | SD | Mean | SD | |
| 0.01× | AFR | Matched | 4789 | 0 | 178 | 72 | 20 | 5 | 6 | 0 | 4 |
| 0.01× | AMR | Matched | 4789 | 0 | 207 | 76 | 14 | 4 | 6 | 0 | 4 |
| 0.01× | EAS | Matched | 4789 | 0 | 214 | 76 | 12 | 4 | 6 | 0 | 4 |
| 0.01× | EUR | Matched | 4789 | 0 | 208 | 76 | 14 | 4 | 6 | 0 | 4 |
| 0.01× | SAS | Matched | 4789 | 0 | 206 | 76 | 14 | 4 | 6 | 0 | 4 |
| 0.01× | AFR | Mismatched | 4789 | 0 | 203 | 75 | 15 | 4 | 6 | 0 | 4 |
| 0.01× | AMR | Mismatched | 4789 | 0 | 202 | 75 | 15 | 5 | 6 | 0 | 4 |
| 0.01× | EAS | Mismatched | 4789 | 0 | 201 | 75 | 15 | 5 | 6 | 0 | 4 |
| 0.01× | EUR | Mismatched | 4789 | 0 | 197 | 75 | 16 | 5 | 6 | 0 | 4 |
| 0.01× | SAS | Mismatched | 4789 | 0 | 201 | 75 | 15 | 5 | 6 | 0 | 4 |
| 0.05× | AFR | Matched | 4789 | 0 | 127 | 49 | 34 | 8 | 19 | 1 | 8 |
| 0.05× | AMR | Matched | 4789 | 0 | 164 | 62 | 26 | 6 | 19 | 1 | 8 |
| 0.05× | EAS | Matched | 4789 | 0 | 176 | 66 | 23 | 6 | 19 | 1 | 8 |
| 0.05× | EUR | Matched | 4789 | 0 | 166 | 63 | 25 | 6 | 19 | 1 | 8 |
| 0.05× | SAS | Matched | 4789 | 0 | 163 | 63 | 26 | 6 | 19 | 1 | 8 |
| 0.05× | AFR | Mismatched | 4789 | 0 | 157 | 60 | 27 | 7 | 19 | 1 | 8 |
| 0.05× | AMR | Mismatched | 4789 | 0 | 159 | 61 | 27 | 7 | 19 | 1 | 8 |
| 0.05× | EAS | Mismatched | 4789 | 0 | 163 | 62 | 26 | 6 | 19 | 1 | 8 |
| 0.05× | EUR | Mismatched | 4789 | 0 | 161 | 61 | 26 | 7 | 19 | 1 | 8 |
| 0.05× | SAS | Mismatched | 4789 | 0 | 160 | 61 | 27 | 7 | 19 | 1 | 8 |
| 0.1× | AFR | Matched | 4789 | 0 | 89 | 35 | 49 | 12 | 26 | 3 | 12 |
| 0.1× | AMR | Matched | 4789 | 0 | 115 | 49 | 38 | 10 | 26 | 2 | 12 |
| 0.1× | EAS | Matched | 4789 | 0 | 126 | 54 | 34 | 9 | 25 | 2 | 12 |
| 0.1× | EUR | Matched | 4789 | 0 | 118 | 50 | 37 | 10 | 26 | 2 | 12 |
| 0.1× | SAS | Matched | 4789 | 0 | 115 | 50 | 38 | 10 | 26 | 2 | 12 |
| 0.1× | AFR | Mismatched | 4789 | 0 | 112 | 48 | 39 | 10 | 26 | 2 | 12 |
| 0.1× | AMR | Mismatched | 4789 | 0 | 113 | 48 | 39 | 10 | 26 | 2 | 12 |
| 0.1× | EAS | Mismatched | 4789 | 0 | 112 | 47 | 39 | 10 | 26 | 2 | 12 |
| 0.1× | EUR | Mismatched | 4789 | 0 | 113 | 48 | 39 | 10 | 26 | 2 | 12 |
| 0.1× | SAS | Mismatched | 4789 | 0 | 112 | 48 | 39 | 10 | 26 | 2 | 12 |

Table S3 Statistics of the simulated admixed triploids, which were used for generating the balanced ROC curve in Fig. 3c. The table includes information about bin sizes, GW sizes, number of GWs per bin and number or reads per GW. All simulated sequences consist of reads 36 bp in length.

| Depth | Ancestry | Reference Panel | Bin size (kbp) | | GW size (kbp) | | GW per Bin | | Reads per GW | | Sampled Reads |
|---|---|---|---|---|---|---|---|---|---|---|---|
| | | | Mean | SD | Mean | SD | Mean | SD | Mean | SD | |
| 0.01× | AFR & EUR | Matched | 4789 | 0 | 187 | 74 | 18 | 5 | 6 | 0 | 4 |
| 0.01× | EAS & EUR | Matched | 4789 | 0 | 207 | 76 | 14 | 4 | 6 | 0 | 4 |
| 0.01× | SAS & EUR | Matched | 4789 | 0 | 206 | 76 | 14 | 4 | 6 | 0 | 4 |
| 0.01× | EAS & SAS | Matched | 4789 | 0 | 208 | 76 | 14 | 4 | 6 | 0 | 4 |
| 0.01× | AFR & EUR | Mismatched | 4789 | 0 | 193 | 74 | 17 | 5 | 6 | 0 | 4 |
| 0.01× | EAS & EUR | Mismatched | 4789 | 0 | 211 | 76 | 13 | 4 | 6 | 0 | 4 |
| 0.01× | SAS & EUR | Mismatched | 4789 | 0 | 207 | 76 | 14 | 4 | 6 | 0 | 4 |
| 0.01× | EAS & SAS | Mismatched | 4789 | 0 | 210 | 76 | 13 | 4 | 6 | 0 | 4 |
| 0.05× | AFR & EUR | Matched | 4789 | 0 | 137 | 53 | 32 | 8 | 19 | 1 | 8 |
| 0.05× | EAS & EUR | Matched | 4789 | 0 | 165 | 63 | 25 | 6 | 19 | 1 | 8 |
| 0.05× | SAS & EUR | Matched | 4789 | 0 | 164 | 62 | 26 | 6 | 19 | 1 | 8 |
| 0.05× | EAS & SAS | Matched | 4789 | 0 | 166 | 63 | 25 | 6 | 19 | 1 | 8 |
| 0.05× | AFR & EUR | Mismatched | 4789 | 0 | 147 | 56 | 30 | 7 | 19 | 1 | 8 |
| 0.05× | EAS & EUR | Mismatched | 4789 | 0 | 171 | 64 | 24 | 6 | 19 | 1 | 8 |
| 0.05× | SAS & EUR | Mismatched | 4789 | 0 | 165 | 63 | 25 | 6 | 19 | 1 | 8 |
| 0.05× | EAS & SAS | Mismatched | 4789 | 0 | 170 | 64 | 24 | 6 | 19 | 1 | 8 |
| 0.1× | AFR & EUR | Matched | 4789 | 0 | 96 | 39 | 45 | 11 | 26 | 2 | 12 |
| 0.1× | EAS & EUR | Matched | 4789 | 0 | 117 | 50 | 37 | 10 | 26 | 2 | 12 |
| 0.1× | SAS & EUR | Matched | 4789 | 0 | 115 | 50 | 37 | 10 | 26 | 2 | 12 |
| 0.1× | EAS & SAS | Matched | 4789 | 0 | 118 | 51 | 37 | 9 | 26 | 2 | 12 |
| 0.1× | AFR & EUR | Mismatched | 4789 | 0 | 104 | 43 | 43 | 11 | 26 | 2 | 12 |
| 0.1× | EAS & EUR | Mismatched | 4789 | 0 | 122 | 52 | 36 | 9 | 26 | 2 | 12 |
| 0.1× | SAS & EUR | Mismatched | 4789 | 0 | 116 | 50 | 37 | 10 | 26 | 2 | 12 |
| 0.1× | EAS & SAS | Mismatched | 4789 | 0 | 120 | 52 | 36 | 9 | 26 | 2 | 12 |

Table S4 Age distribution of the patients that went through an IVF process at the Zouves fertility center between 2015 to 2020.

| Age group | Percentage |
|---|---|
| 20 – 25 | 0.1% |
| 25 – 30 | 1.9% |
| 30 – 35 | 19.2% |
| 35 – 40 | 40.2% |
| 40 – 45 | 32.9% |
| 45 – 50 | 5.1% |
| 50 – 55 | 0.6% |
| 55 – 60 | 0.1% |
| 60 – 65 | 0.0% |

### Supplemental Figures

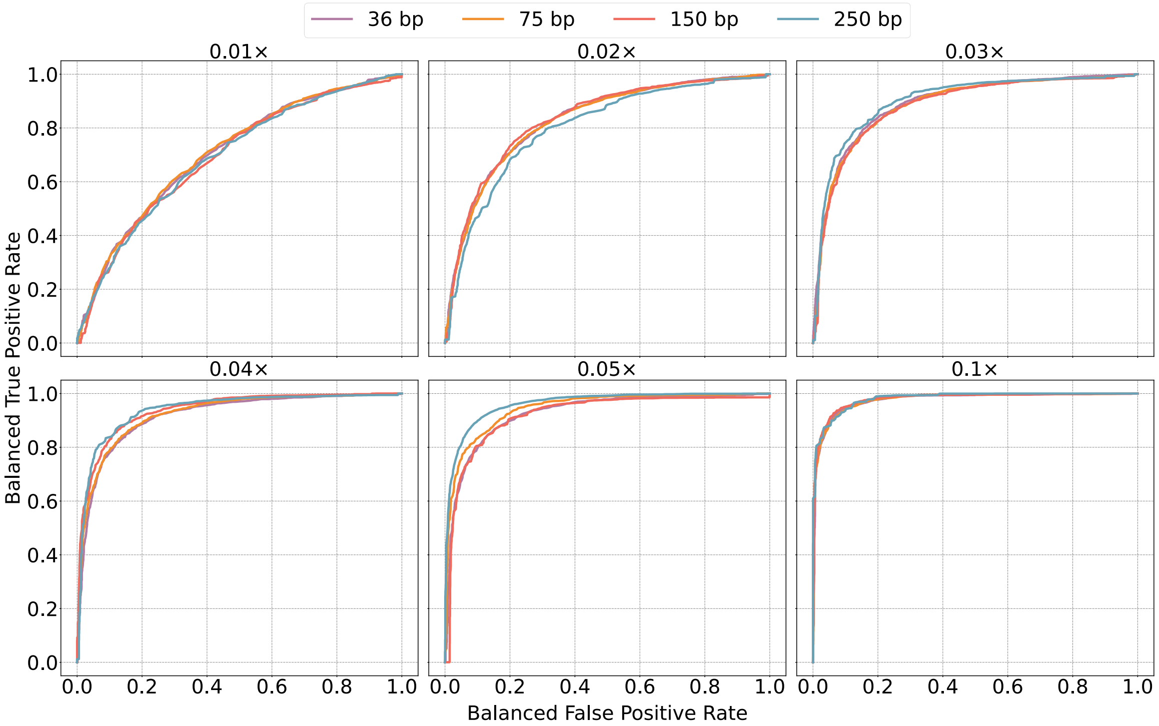


Figure S1 Balanced ROC curves for BPH vs. SPH with matched reference panels, varying depths of coverage and read length. Each balanced ROC curve reflects an average over 10 bins across Chromosome 21. We averaged both the BTPR and BFPR for common $z$-scores across bins. The plots show that at extremely low coverages, short reads perform better because they provide more uniform coverage. As the coverage increases and uniform coverage is achieved also with long reads, the situation changes. Long reads are more likely to to overlap with more than one SNP and therefore allow us to distinguish between the different haplotypes with greater accuracy.

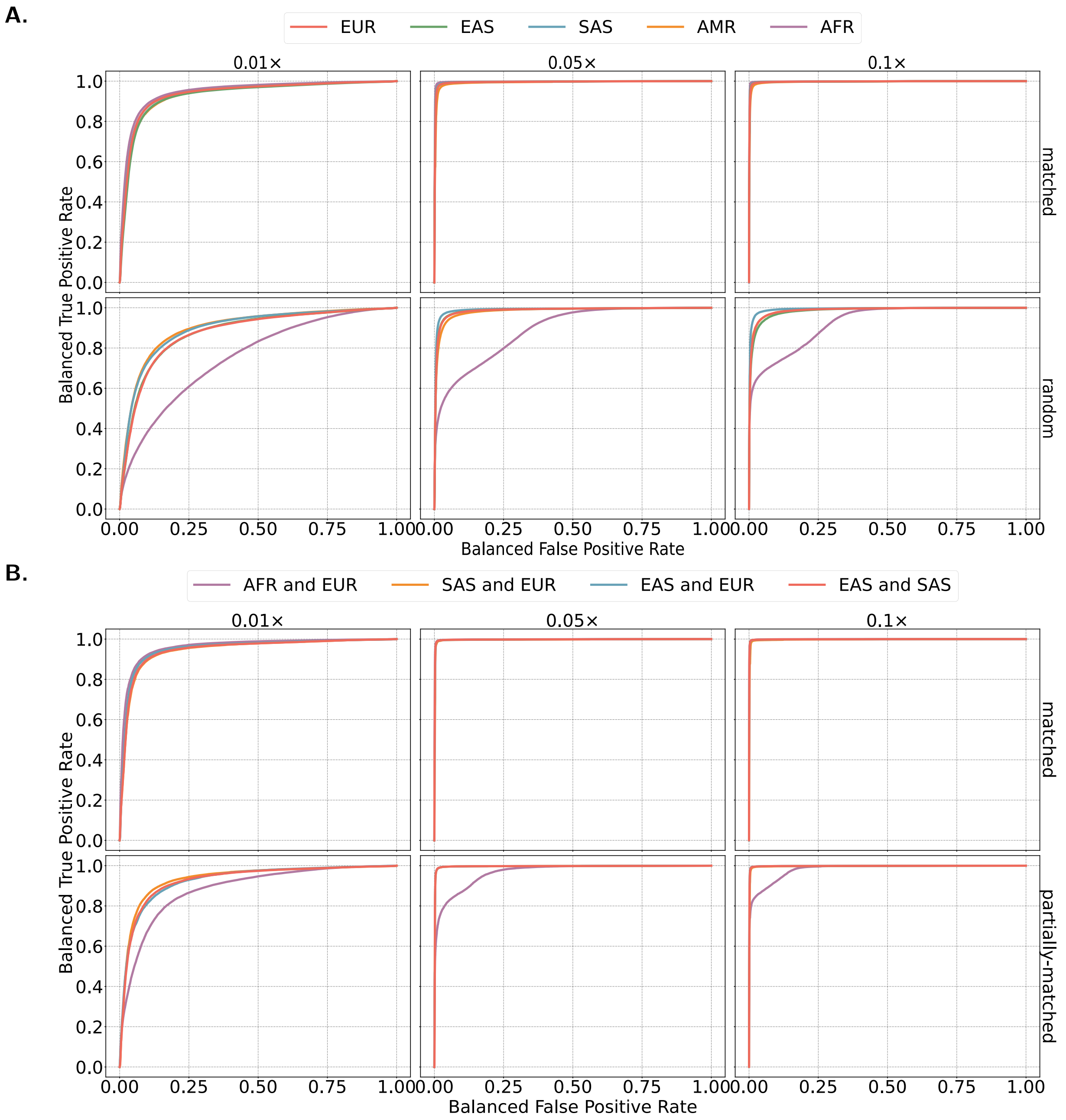


Figure S2 A. Balanced ROC curves for monosomy vs. disomy with matched and random reference panels of non-admixed embryos, varying depths of coverage. B. Balanced ROC curves for monosomy vs. disomy with matched and partially-matched reference panels of admixed embryos, varying depths of coverage. Here the randomly mismatched reference panel corresponds to either the paternal or maternal ancestry. Each balanced ROC curve reflects an average over bins across the genome. We averaged both the BTPR and BFPR for common $z$-scores across bins.

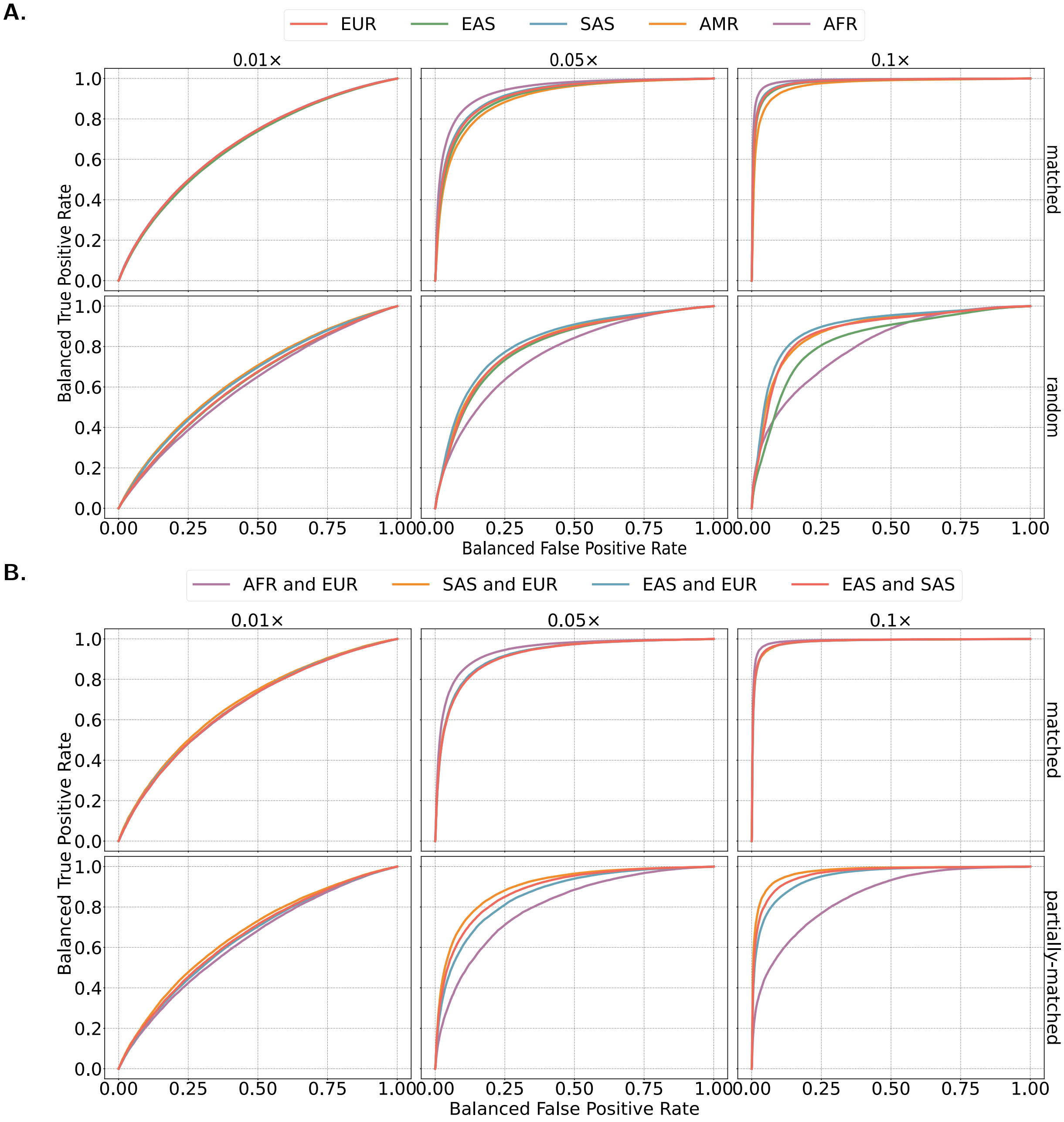


Figure S3 A. Balanced ROC curves for BPH vs. disomy with matched and random reference panels of non-admixed embryos, varying depths of coverage. B. Balanced ROC curves for BPH vs. disomy with matched and partially-matched reference panels of admixed embryos, varying depths of coverage. Here the randomly mismatched reference panel corresponds to either the paternal or maternal ancestry. Each balanced ROC curve reflects an average over bins across the genome. We averaged both the BTPR and BFPR for common $z$-scores across bins.

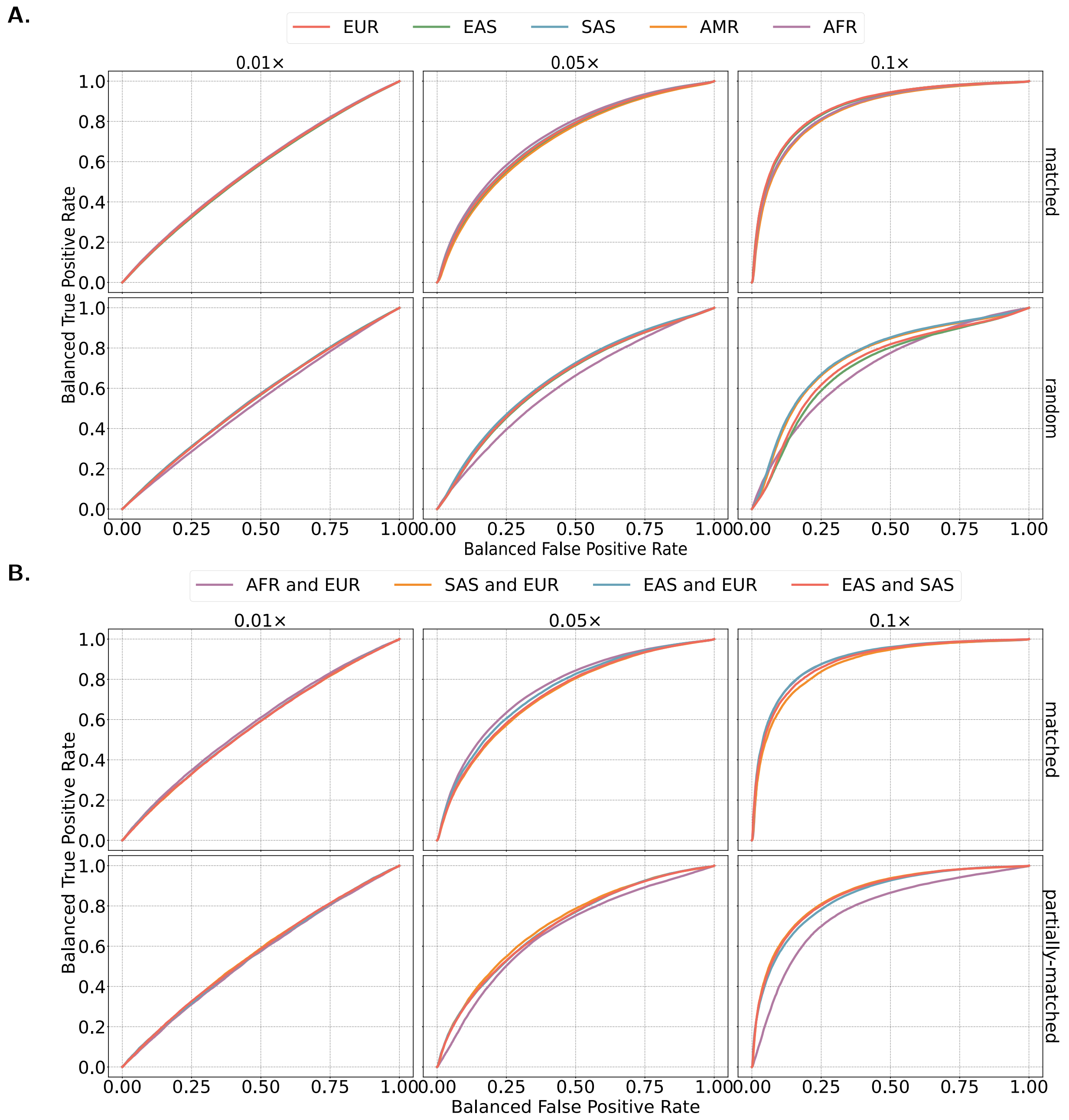


Figure S4 A. Balanced ROC curves for SPH vs. disomy with matched and random reference panels of non-admixed embryos, varying depths of coverage. B. Balanced ROC curves for SPH vs. disomy with matched and partially-matched reference panels of admixed embryos, varying depths of coverage. Here the randomly mismatched reference panel corresponds to either the paternal or maternal ancestry. Each balanced ROC curve reflects an average over bins across the genome. We averaged both the BTPR and BFPR for common $z$-scores across bins.

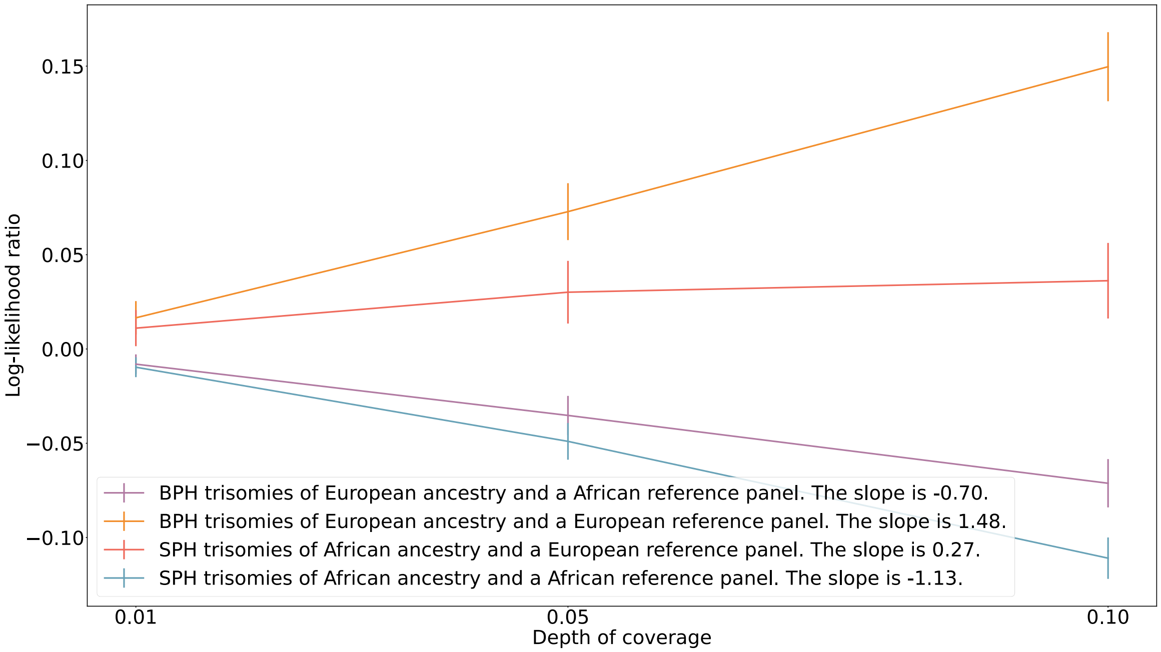


Figure S5 The LLR of BPH over SPH is averaged over many simulated trisomies. BPH and SPH trisomies were simulated for depths of 0.01×, 0.05× and 0.1×. The plot demonstrates how misspecification involving a reference panel with different rates of heterozygosity than the target sample affects the LLR per genomic window. We estimated the slopes of each line using the least squares method.

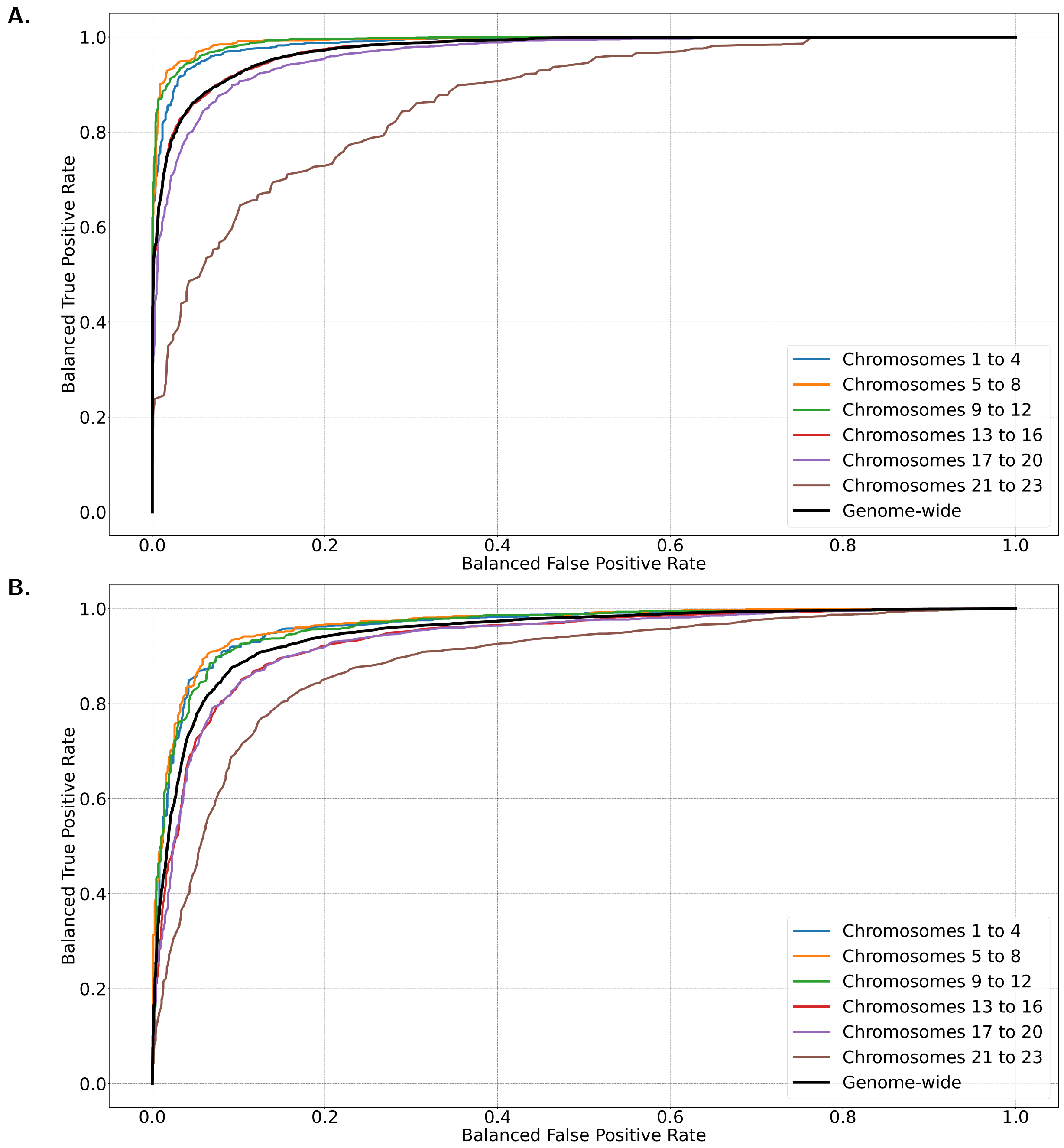


Figure S6 A. Balanced ROC curves for BPH vs. SPH with matched reference panels and depths of coverage matched to those observed within the Zouves dataset. Each balanced ROC curve reflects an average over bins across the genome. We averaged both the BTPR and BFPR for common $z$-scores across bins. B. Balanced ROC curves for disomy vs. monosomy with matched reference panels and depths of coverage matched to those observed within the Zouves dataset. Each balanced ROC curve reflects an average over bins across the genome. We averaged both the BTPR and BFPR for common $z$-scores across bins.

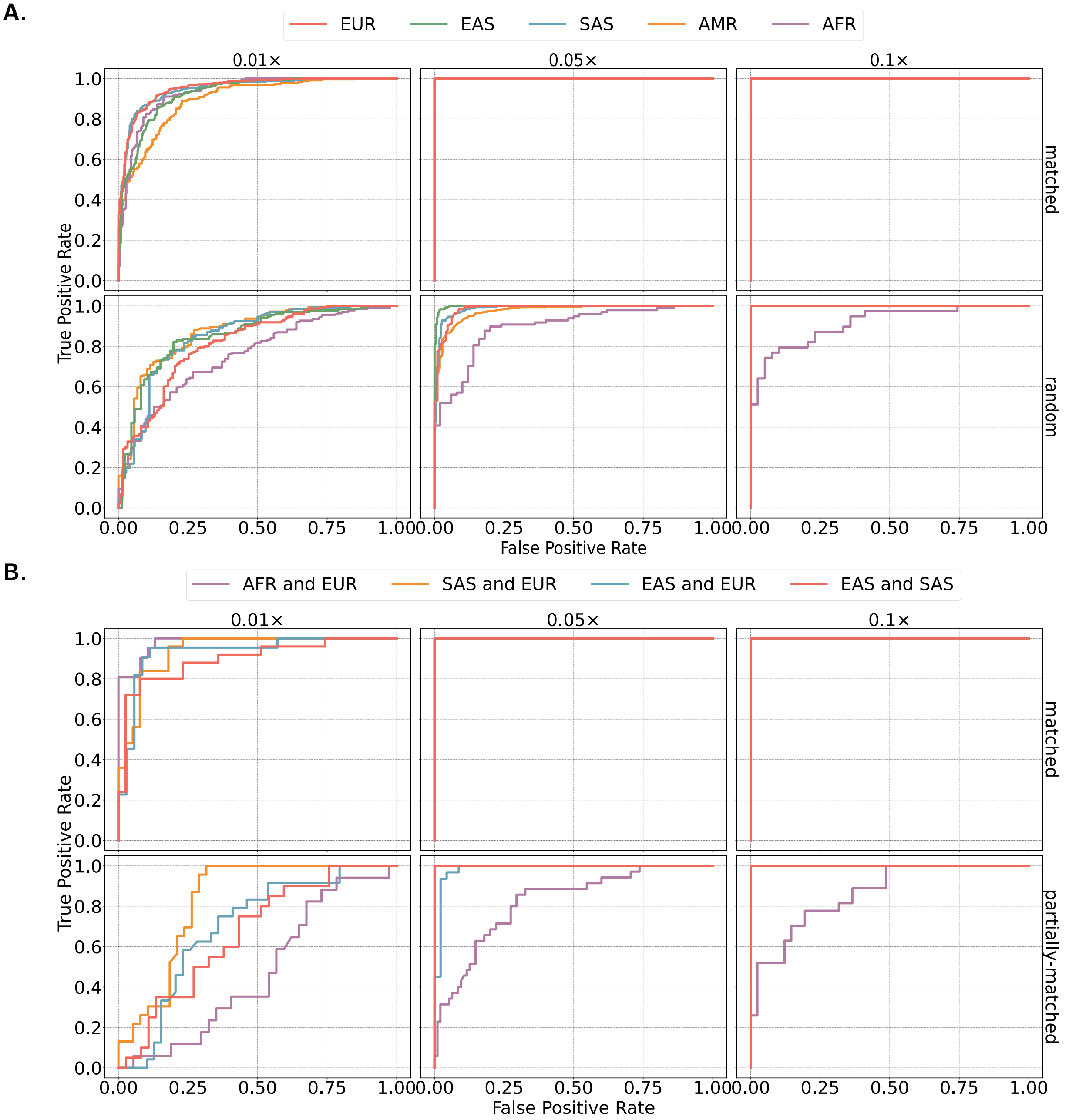


Figure S7 A. Standard ROC curves of triploidy vs. diploidy for non-admixed embryos with matched and random reference panels and varying depths of coverage. B. Standard ROC curves of triploidy vs. diploidy for admixed embryos with matched and partially-matched reference panels and varying depths of coverage. Here the randomly mismatched reference panel corresponds to either the paternal or maternal ancestry. TPR and FPR denote true and false classification of diploidy, respectively. Each trisomic chromosome of the triploid embryo comprises a mixture of BPH and SPH tracts, with details of these simulations described in the Methods section.

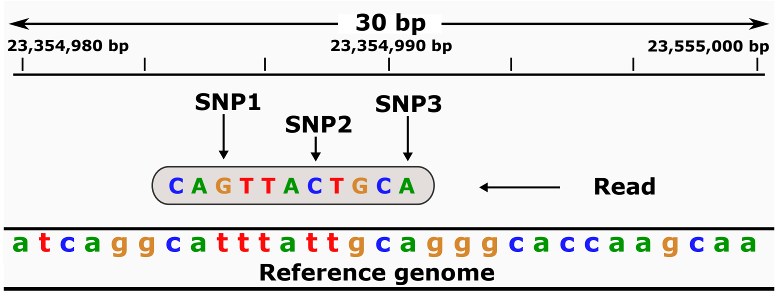


| Haplotype | Joint Frequency | Score Increment $= \begin{cases} 1, & 0.1 < f < 0.9 \\ 0, & \text{otherwise} \end{cases}$ |
|---|---|---|
| $GTC$ | 0.22 | +1 |
| $GTA$ | 0.02 | 0 |
| $GCA$ | 0.15 | +1 |
| $GCC$ | 0.28 | 0 |
| $TTC$ | 0.01 | 0 |
| $TTA$ | 0.37 | +1 |
| $TCA$ | 0.03 | 0 |
| $TCC$ | 0.01 | 0 |
| | | Total score: 3 |

Figure S8 (Top) A read that is aligned to a reference genome. (Bottom) A table of possible haplotypes in the chromosomal region that overlaps with the read, the frequency of the haplotypes (as estimated from a reference panel) and the score of the haplotype.

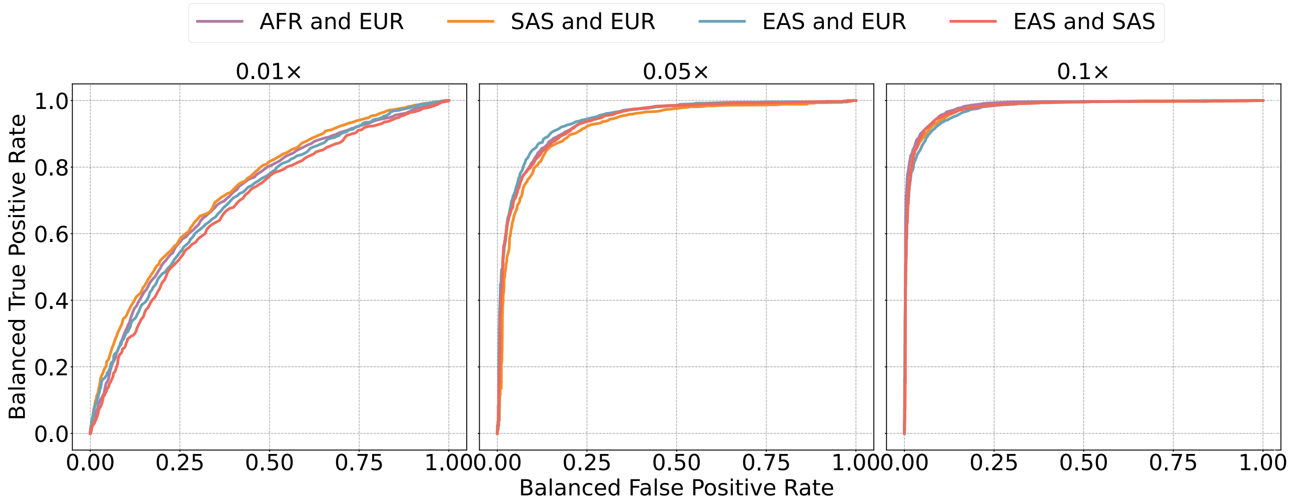


Figure S9 Balanced ROC curves for BPH vs. SPH for distant and arbitrary admixtures, based on simulating Trisomy 21 with varying depths of coverage. The ancestry proportions for each simulated admixture scenario is 50% for each of the two respective populations. More specifically, we partitioned chromosome 21 into 40 equally sized intervals. For each region, three haplotypes were drawn with equal probability of drawing each from either of the two populations. Each balanced ROC curve reflects an average over bins across Chromosome 21. We averaged both the BTPR and BFPR for common $z$-scores across bins.
